# Sound-evoked auditory neurophysiological signals are a window into prodromal functional differences in a preclinical model of Alzheimer’s Disease

**DOI:** 10.1101/2025.08.07.669134

**Authors:** Aysegul Gungor Aydin, Pranav Manoj, Faiza Ramadan, Sarah Rajan, Elias Youssef, Elizabeth B. Torres, Kasia M. Bieszczad

## Abstract

Hearing is the largest modifiable mid-life risk factor for Alzheimer’s Disease (AD), yet its link to dementia remains unclear. We identified a neurophysiological biomarker of AD risk using the non-invasive, rapidly acquired, and clinically translatable auditory brainstem response (ABR) in normal hearing knock-in rats (Swedish familial AD risk variant to Amyloid precursor protein, App^S^; male and female). Human ABRs have been proposed as a biomarker for AD and related dementias. The novel metric reported here is derived from multidimensional parametric feature extraction on the distribution statistics of repeated single-trial ABR traces. We report accurate prediction of genetic risk for AD risk in young and aged rats: App^S^ separate clearly from healthy humanized (App^H^) in sex- and age-dependent manners. Notably, auditory learning during young adulthood shifted the App^S^ ABR signature towards a healthy App^H^-like state that maintained over time into older age. Altogether the findings support the utility of the ABR to track disease state, progression, and effects of intervention, and point to a central neural generator of auditory dysfunction related to AD risk. ABRs could provide a very early biomarker for detection of AD risk and used to test the synergy of auditory and cognitive functions in human dementia.

## INTRODUCTION

Hearing loss is the greatest modifiable risk factor in midlife for developing Alzheimer’s Disease (AD) and related dementias (ADRD)^1–3^. Although the significance of auditory system processes in dementia has been validated by 20+ years of research showing links between hearing loss, cognitive decline, and cortical atrophy, mechanisms to link hearing ability with dementia remain unclear^4–7^. Hypotheses that range from causal, indirect, or common cause mechanisms to link hearing ability with dementia diagnosis are now being actively investigated in multiple disciplines^8^ as a timely response to emerging evidence that correcting hearing loss with hearing aids in humans does not always mitigate AD risk^9,10^. The relationship may instead involve functional integration of auditory cortical and subcortical circuitry that directly affects learning and memory processes, the first cognitive abilities affected in ADRD. For example, animals and humans that successfully learn and remember have neuroplasticity events that rapidly and stably reorganize auditory cortical circuits to strengthen representations evoked by behaviorally salient sounds^11–17^. Neurophysiological failures in reliable and consistent neural transmission will make it harder to hear—or rather, to listen and understand—sound cues^18,19^. Indeed, auditory function may be particularly susceptible to functional changes in the dynamics of synaptic transmission evident very early in neurodegenerative disease, potentially explaining the link between *mid-life* hearing before pathology emerges and *later* dementia risk^20,21^. This creates a compelling clinical opportunity for earlier AD/ADRD diagnoses and disease monitoring via the auditory system.

A prime auditory neurophysiological candidate to act as a sensitive biomarker of AD risk is the auditory brainstem response (ABR), which can be obtained by non-invasive EEG-like recordings. This “brainstem response” is an apparent misnomer since the ABR reflects integrated function across the entire auditory system that appears as a series of waves (I, II, III, IV, V, etc.) in a typical trial-averaged response. Each sound-evoked ABR wave has an amplitude and latency that reflect population activity from generally-agreed upon ascending central neural generators from the auditory nerve, to subcortical and higher auditory cortical areas^18,22^. While the gross morphology of identifiable waves is similar across subjects, the detailed morphology of the ABR is unique between individuals (in humans and rodents) and is stable over time. Thus, the ABR offers a rapid, reliable, objective, highly reproducible, and non-invasive method to assess the integrity and functioning of central auditory function. It can reflect stimulus-locked, synchronous neural firing from nuclei along the ascending auditory pathway with precise temporal resolution ^23^ and small inter- and intrasubject variability ^24,25^. Although the ABR is widely used clinically to assess hearing thresholds, it effectively captures synaptic transmission along the longitudinal extent of the central auditory pathway, including axonal conduction and synaptic delays between ascending and descending projections of distant auditory nuclei ^26^. By capturing neural transmission over many milliseconds, the ABR confers sensitivity to structural and functional disruptions that might be caused by injury, aging, neuropathy or demyelinating diseases^27–29^.

As such the ABR is a multi-purpose non-invasive tool well-suited for studies investigating central nervous system integrity. We and others have used it as an auditory fingerprint with individual differences gained by a subject’s experiential life history^30^ and potential disease states^29,31,32^. ABR wave amplitude and latency have been suggested as markers of mild cognitive impairment (MCI) and AD^33^, but their clinical impact may be limited in predicting disease state or severity ^34,35^. Additional analytical methods might improve predictive performance beyond conventional ABR measures in clinical populations, highlighting an opportunity to substantially enhance sensitivity and specificity relative to standard measures while using the same data^36^. Here, we apply a novel objective analytical method to functionally characterize ABRs by using a multi-dimensional parametric feature space obtained from single-trial data^37^. This method allows a neural signature to be characterized beyond qualitative ABR morphology, amplitude, latency, or subjective threshold assessments^38^. By applying mathematical modelling to the distributions of single trial sound-evoked response variability across hundreds of repeated sound presentations, the parameter space in which the details of individual subjects’ responses can be classified is expanded, which may increase the sensitivity and specificity of an ABR biometric for disease risk. Similar analytical methods applied to single-trial ABR data from human neonates have successfully identified unique features of microscale variability measures that can be used remarkably early in human life to correctly predict future diagnosis of neurodevelopmental disorders that are not easily recognized until the age when neurotypical children become verbal^31^. Thus, this biomarker has the benefit of very early detection, prodromal to the disease.

In this report, we initially present a novel analytical method applied to the ABR as a diagnostic tool to characterize very early neurophysiological dysfunction with AD/ADRD disease risk. It is here presented in a preclinical rat model of AD that mimics early-onset (<65 years of age) human Familial AD (FAD) via a CRISPR/Cas9-mediated “Swedish” risk variant knock-in to the genomic amyloid-beta (Aβ) precursor protein (*App*) locus^39–42^. Of the many preclinical models available in rodents, the rat has been the organism of choice for most behavioral, memory, and cognitive research, as the rat has a high degree of intelligence and high level of cognitive ability, including flexible, knowledge-dependent behavioral control and the ability to learn complex experimental contingencies, which has inspired an increase in the use of rats to model human disease. Moreover, rat and human APP differ by only 3 amino-acids in the Aβ region, and rats have the same tau isoforms as humans (unlike the mouse)^43^ and tend to survive to adulthood with homozygous mutations to AD gene targets that can be lethal in mice (e.g., for *Psen1*)^42,44^. Thus, we used FAD rat models with imposed genetic risk as a valuable opportunity to identify very early signatures of AD prodromal to characteristic tau and/or amyloid pathology that are likely to translate to the human context. It is important to note that the *App*^S^ rat has APP pathogenic processing, and changes in synaptic signaling across the lifespan compared to humanized controls^39–41^, without developing characteristic AD-like pathology until older age or with additional disease mutations^45,46^. Therefore, ABRs from *App*^S^ rats were investigated here to model the mid-life modifiable risk context that links hearing and dementia, which at this life timepoint in humans, occurs *without* classical AD pathology^47^.

Capitalizing on the opportunity to study a preclinical model without overt neuropathology, we sought to determine whether a sound-evoked neurophysiological signal could screen known genetic risk for AD between the *App* Swedish risk variant vs. humanized knock-in rats and track risk severity in aging—possibly independent of Aβ deposition^46,48^. Further, since auditory learning can stably modify neural activity that can be captured by ABR features (Skoe 2013; Rotondo & Bieszczad 2021), we used behavioral sound-reward training as an intervention to test whether the same sound-evoked neurophysiological signal could track the effects of intervention. Comparing conditions with expected differences in APP pathogenic processing opens a unique window to reveal the diagnostic potential of functional synaptic pathological mechanisms of neurodegeneration evident in the auditory nervous system that are prodromal to amyloid or tau pathology, emerging very early in life and evident with access to neurophysiological phenotyping via the ABR.

## RESULTS

Single trials of tone-evoked ABRs were collected (512 repetitions of 5 msec, 5.0 kHz pure tones at 70 dB SPL; 21 Hz repetition rate) in male and female homozygous *App*^S^ or *App*^H^ rats. Click-evoked ABR hearing thresholds were considered normal and not different between genotypes (*App*^S^ hearing threshold: 42.14 ± 1.84 dB SPL vs. *App*^H^ hearing threshold: 45 ± 2.23 dB SPL, t (8.631) = 0.986, p = 0.351; Welch’s *t*-test) (**FIGURE S1A-D & FIGURE S2, SUPPLEMENTARY TABLE S1**). However, they were different between sexes (male hearing thresholds: 50 ± 0 dB SPL vs. female hearing thresholds: 41.11 ± 1.11 dB SPL, t (8) = 8.00, p < 0.0001; Welch’s *t*-test) (**SUPPLEMENTARY FIGURE S1B,E**). Thresholds increased slightly with aging, as expected (5 months old: 40 ± 1.29 dB SPL vs. 12+ months old: 46.67 ± 1.67 dB SPL, t(10) = 3.162, p = 0.010; unpaired t-test) (**SUPPLEMENTARY FIGURE S1C,F**). Next, single-trial ABR data were analyzed as a continuous timeseries of “micromovement” peaks, the mathematics of which have been described in detail (see *Materials & Methods, Data Analysis: Single-trial ABRs timeseries analysis* and *Supplemental Material: Pipeline to derive the micro-movement spikes and Gamma representation from positively rectified ABR signals*).

### Rat ABR micromovement spikes can be well characterized by a Gamma process

The normalized micro-peaks comprising the micro-movement spikes of rat ABR signals, taken trial by trial instead of averaging the waveforms, follow the continuous family of probability distributions. The Gamma mean and Gamma variance can be used to compute the Gamma noise to signal ratio, giving us a sense of individualized dispersion of the response fluctuations around the local mean response. Using a time window of 10 ms that transformed ABR data obtained at an initial sampling resolution of 195200 Hz and filtered between 0.3 and 3.0 kHz, we captured the stochastic shifts of the responses and distinguished different classes of animals accordingly. Furthermore, we characterized the individual stochastic shifts to appreciate the outcomes from different stimulus exposure. The shape and scale parameters of the continuous Gamma family of probability distribution functions can be mapped as points on the Gamma parameter plane with 95% confidence intervals. This parameter plane is spanned by the shape and scale (dispersion) of the empirically estimated distribution family. The log-log scatter of points on the Gamma parameter plane followed a tight linear fit such that as the scale parameter decreased, the shape parameter increased. The shape parameter at 1 is the special case of memoryless exponential distribution, while higher values represent heavy - tailed distributions, and even larger values (> 100) fall along the symmetric, Gaussian-like ranges of the Gamma family. Finding that empirical data follows this distribution family, rather than a priori adopting a theoretical distribution, has implications for translational rat models in general. The nuanced data offered by this new empirical approach captures stochastic patterns of neurodevelopment or neurodegeneration analogous to those previously quantified in humans using this model (**FIGURE 1**)^31,49^.

**Figure 1:**
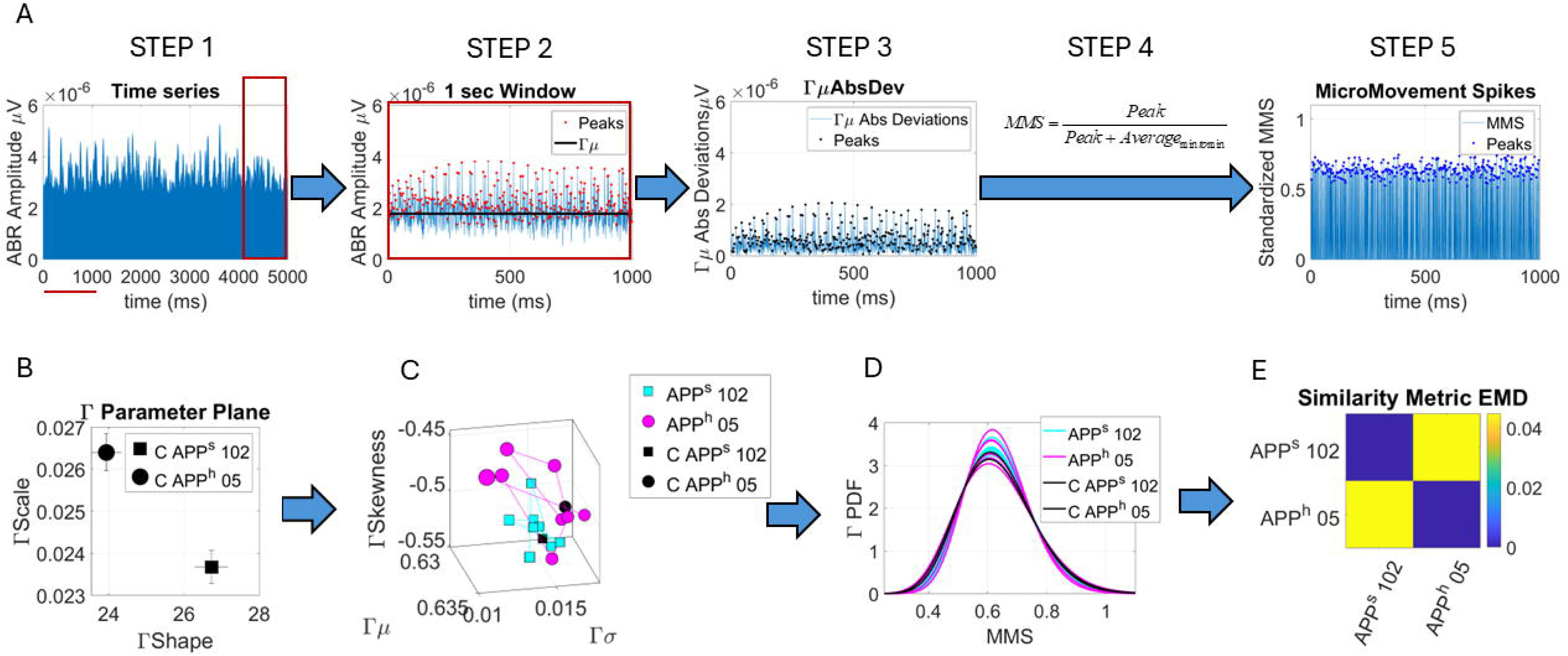
Data standardization and analytical pipeline. **(A)** Step 1 provides the raw time series data 5 seconds long expressed in milliseconds. One-second long sliding window with 50% overlap is used to traverse the data. Red rectangle shows a representative window zoomed in step 2. Step 2 shows the peaks of the data which are gathered into a frequency histogram and subject to maximum likelihood estimation. The continuous Gamma family of probability distribution functions is the best fit and the first moment, the mean value thus obtained is plotted as a black line. The absolute deviations from the mean are obtained and shown in step 3 along with the peaks which will be subject to standardization by equation in Step 4. This normalization scales out allometric effects due to anatomical disparities across animals, such that we can appropriately compare outcomes across different animals. Step 5 provides the micro-movement spikes MMS, waveforms used in the stochastic analyses. The MMS are thus standardized deviations from the empirically estimated mean. At zero value, they are the mean activity. **(B)** The MMS provide the micropeaks to empirically estimate the family of distributions best fitting the data in an MLE sense. The continuous Gamma family of probability distribution functions provides the best MLE estimates which are plotted with 95% confidence intervals on the Gamma parameter plane spanned by the Gamma shape and Gamma scale (also the Noise to Signal Ratio, NSR). **(C)** the full stochastic trajectory from the sliding window yields the stochastic range of each animal, plotted here along with the centroids of this range in B. **(D)** The family of Gamma PDFs shows the non-stationary nature of the data and the non-normality (heavy right tails). **(E)** This is further quantified by pairwise comparison of the PDFs using the Earth Mover’s Distance, an appropriate similarity metric to compare PDFs as points of a probability space. A*bbreviations*: MMS, micromovement spikes; PDF, probability distribution function.

### ABR micromovements are sensitive to FAD-risk in rats

Aged *App*^S^ rats (>12 months old) clearly separate within sex from controls *App*^H^, as their responses feature different empirical probability distribution functions. This is appreciated through their representation as points of the Gamma parameter plane with non-overlapping 95% confidence intervals (**FIGURE 2**). Proper similarity metric to assess distance in probability space also captures the differences along with the two-distribution Kolmogorov-Smirnov test comparing the unfolded distribution pairwise (p<0.01; **SUPPLEMENTARY FIGURES S3-S5**). Separation is remarkably still evident in an independent cohort of younger animals (5 months old) (only partial overlap in 95% confidence intervals; and less clear genotype-specific dissimilarity; **FIGURE 3**; **SUPPLEMENTARY FIGURES S6, S7**). Notably, the empirical distributions from the responses of aged animals separate further in the same Gamma parameter plane, consistent with the interpretation that thus obtained ABR metrics capture progressive dysfunction with AD risk, e.g., aging-dependent neurodegeneration. Interestingly, we found that the same amplitude-dependent parameter dimensions did not separate animals with an alternate risk allele, the LF autosomal dominant risk variant to presenilin (*Psen1)* that encodes a γ-secretase component linked to familial AD (**SUPPLEMENTARY FIGURE S8**), which suggests that functional differences could be specific to risk genotype. Thus, nuanced trial-by-trial ABR data can separate AD risk and progression. Differentiation may be dependent on neurophysiological characteristics captured by the individualized and properly standardized ABR signal likely reflecting specific features of disease-etiology.

**Figure 2:**
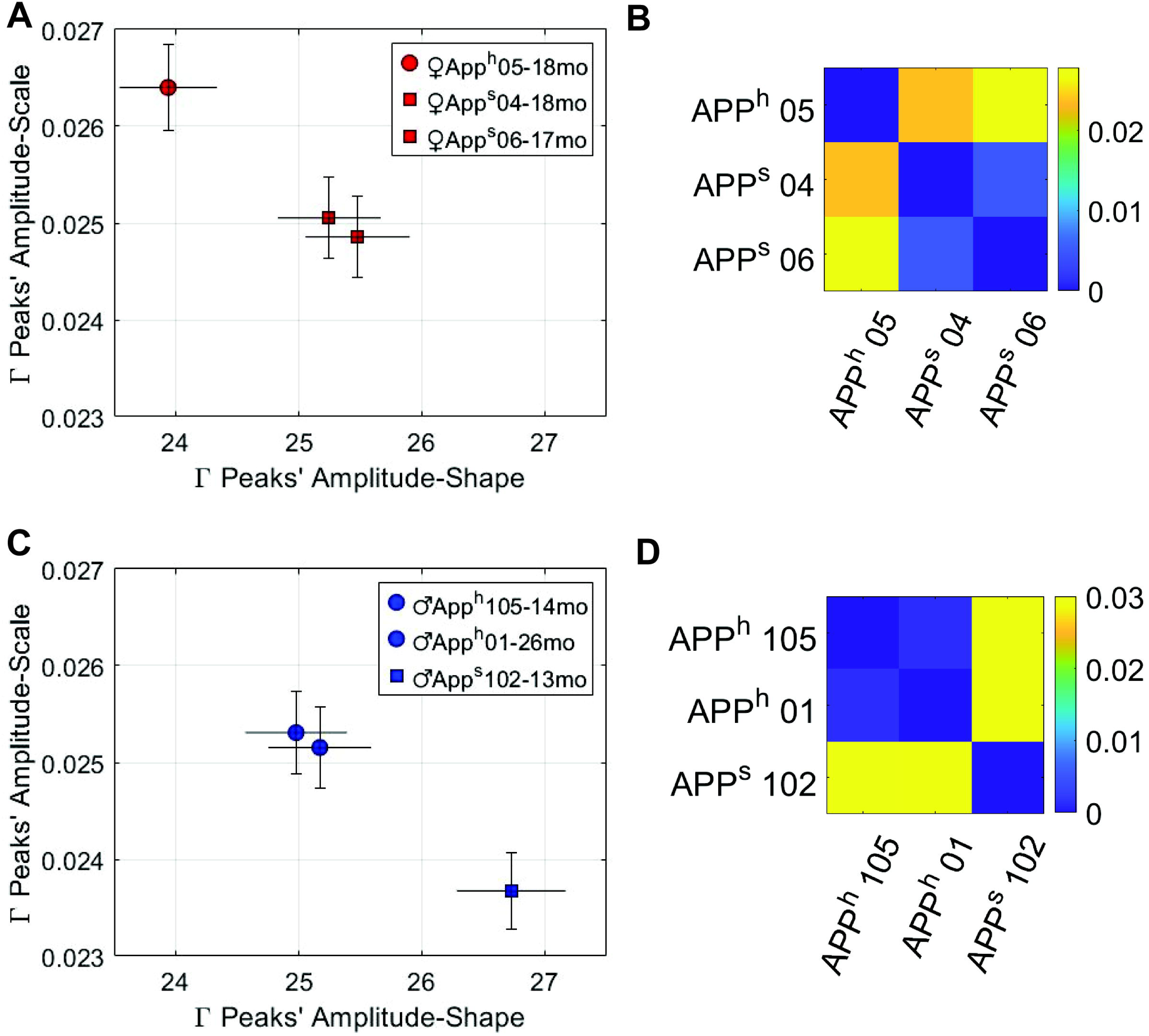
Multivariate time-series analysis of *auditory brainstem response (ABR)* data predicts AD risk in aged rats. ABR response characteristics distinguish AD risk mutant *App^S^* rats (*App* Swedish homozygous, square icons) from *App^H^*controls (*App* humanized allele, circle icons) in a 2-dimensional state space defined by micromovement (gamma), shape (x-axis), and **scale** (y-axis). For (**A**) shows separation in the peaks’ amplitude in the ABR waveform for females, and (C) shows separation for males. Each symbol represents an individual subject, and error bars indicate within-trial fluctuations in the peak amplitude, shown as 95% confidence intervals (crosshairs). Red denotes females and blue denotes males. There is no overlap between risk vs. wildtype genotypes in both sexes. Similarity between distributions in the Gamma parameter space was quantified using Earth Mover’s Distance (EMD) and visualized as pairwise colormaps (B, D), where cooler colors indicate greater similarity and warmer colors indicate greater dissimilarity. App 04 and 06 mutants exhibit greater similarity to each other and are more distinct from App 05 humanized (B). Similarly, in males, App 105 and App 01 humanized show greater similarity to each other and are more distinct from App 102 mutant. Legend shows genotypes. Abbreviations: *App*, amyloid precursor protein.

**Figure 3:**
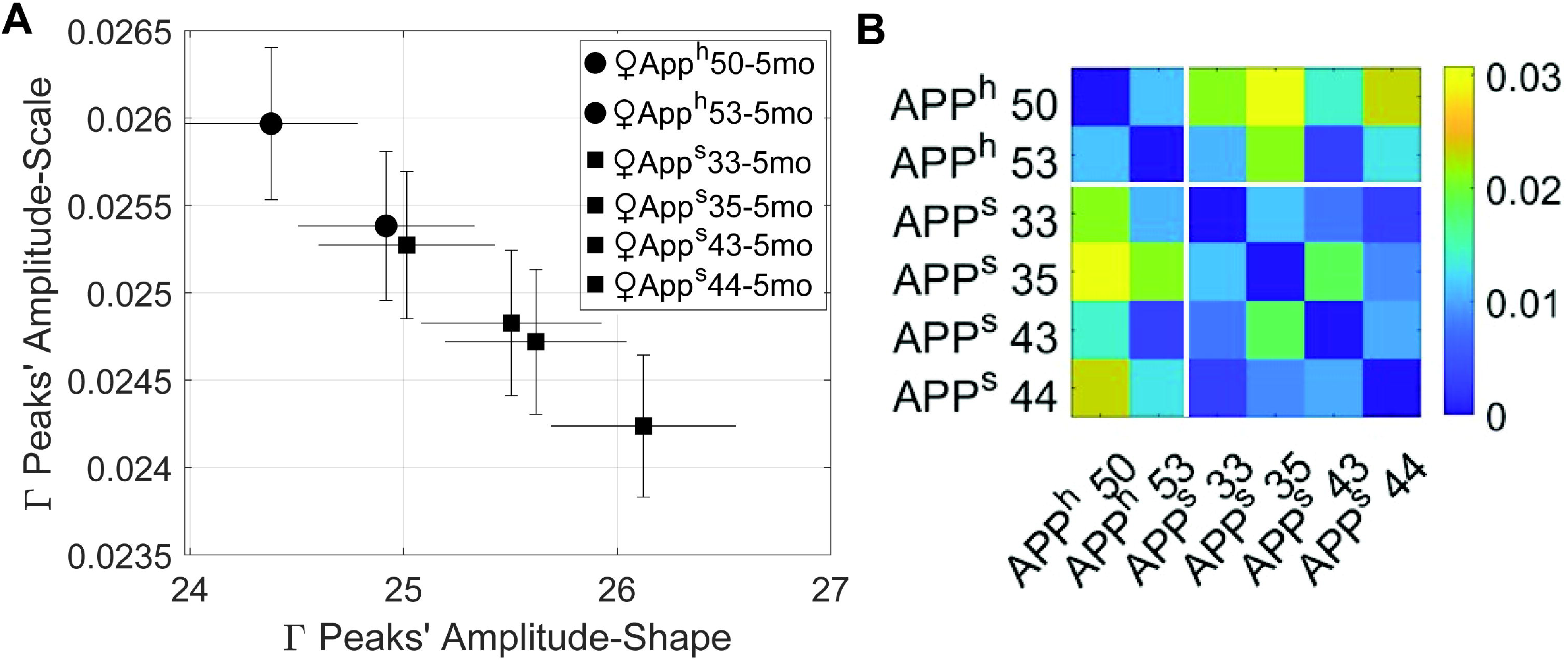
*Auditory brainstem response (ABR)* waveform features capture AD risk in young (5-month-old) female rats. (A) Same as in Fig. 2, but in young age, fluctuations in the peaks’ amplitude in the ABR distinguish AD risk mutant female *App^S^*rats (square icons) vs *App^H^*controls (circle icons). Only one *App^S^* rat fails to distinguish from the control subject’s confidence space, consistent with individual differences in susceptibilities to genetically driven phenotypes. Otherwise, there is a robust separation between AD risk and wildtype genotypes. Points represent the gamma shape and scale parameter space to localize individual subjects with 95% confidence intervals. (B) In contrast to aged animals, genotype-dependent similarities between distributions in the Gamma parameter space are weaker than those observed in aged animals.

### ABR micromovements shift after a bout of auditory-cognitive training

Tracked longitudinally, we report that the ABR of *App*^S^ rats can transform with sound-guided cognitive training to overlap with *App*^H^ (**FIGURE 4**; **SUPPLMENTARY FIGURE S9**). Remarkably, 2-3 weeks of directed auditory training in a simple tone-reward task (**SUPPLMENTARY FIGURE S10**) ameliorates genotype-dependent ABR differences in a way that appears to shift auditory responses to become indistinguishable from controls. The shift persists for at least 9 months (as long as recordings were possible). An initial analysis from the same data shows no differences in hearing thresholds with training (pre-training: 56.67 ± 3.33 dB SPL vs. post-training: 58.33 ± 4.01 dB SPL, t (5)= 1.00, p = 0.363; paired t-test; **SUPPLEMENTARY TABLE S1)**, yet pre- vs. post-training ABRs from *App*^S^ vs *App*^H^ trained rats show an increased amplitude of the second peak wave (PW2) of the ABR (**SUPPLEMENTARY FIGURE S11**; **SUPPLEMENTARY TABLES S2, S3**). The relative increase in PW2 amplitude scales with the success of tone-reward memory formation measured behaviorally (Pearson’s correlation, two-tailed: R = 0.790; p = 0.0197) (**SUPPLEMENTARY FIGURE S11C)**.

**Figure 4:**
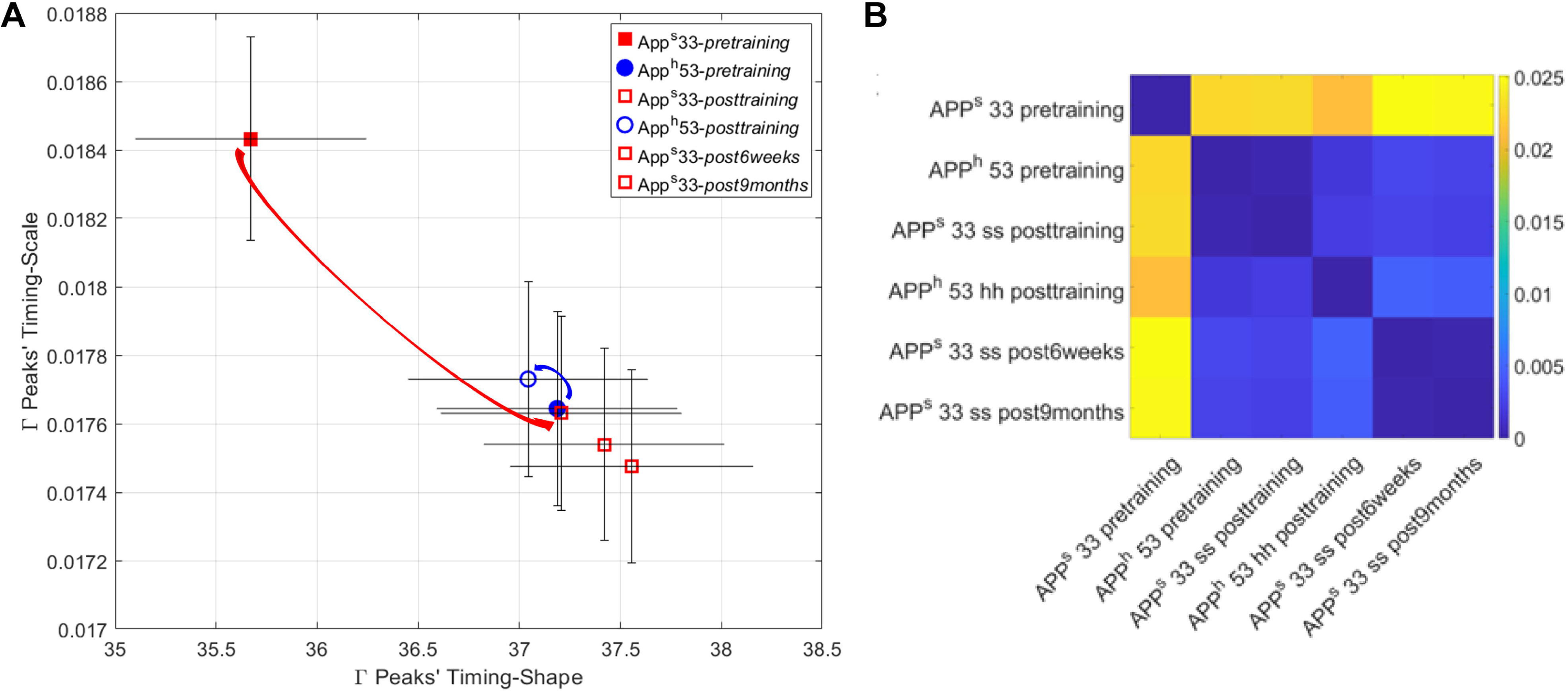
Auditory training shifts *auditory brainstem response (ABR)* waveform dynamics in *App^S^* and *App^H^* rats and shifts the auditory responses of *App^S^* rats to become indistinguishable from *App^H^* rats. (A) As before, stochastic patterns of multivariate characteristics in the ABR distinguish AD risk mutant *App^S^* rats (filled red square) from *App^H^* controls (filled blue circle) in a 2-dimensional state space (gamma shape and scale), which is evident before training (filled blue circle). However, following training in a simple sound-reward go/no-go learning task, there is a functional change that is read out in our ABR metric (open red circles). Remarkably, the APP^s^ ABR is shifted to the *App^H^* condition (open and filled blue circles) with this intervention. Curved arrows indicate the direction of change in ABR state space from pre-training (filled symbols) to post-training and follow-up time points (open symbols). Red curved arrow shows effect in this space state towards the overlapping areas of parameter space. This effect on *App^S^* lasts at least up to 9 months (open red circles show immediate, 6 weeks and 9 months post-training). Blue curved arrow indicates trajectories of change for the *App^H^* rat. Error bounds show the variability in the relevant parameter of an individual subject’s profile in this state space over time as a 95% confidence interval (crosshairs). Data from two female rats are shown, using longitudinal recording of ABRs from each to characterize dramatic shifts induced by training intervention. (B) *App^S^* and *App^H^* rats in post training conditions show greater similarity to each other and strikingly distinct from *App^S^* in pre-training condition in the distances in Gamma probability space.

## DISCUSSION

We report three major findings: First, the rat ABR micromovement spikes, away from the animal’s empirical response mean, and properly normalized to scale out allometric effects from animals’ anatomical disparities across the cohort, are best fit by the continuous Gamma family of probability distributions in a maximum likelihood estimation sense. Accordingly, the Gamma shape and scale parameters derived from the standardized micro-movement spikes of trial-by-trial ABR can be used to represent individual animals and cohorts of animals on the Gamma parameter plane with 95% confidence. As in humans, the log-log representation follows a tight linear relation whereby as the Gamma scale parameter decreases, the shape parameter increases towards the Gaussian ranges of the Gamma family. The range of distribution parameters of this family thus can be used to characterize the distribution ranges for each animal and those of different classes of animals, including as they shift with aging and intervention on properly defined parameter spaces. In simplest terms, this finding shows a promising case of reverse translation opportune for bridging clinical studies to preclinical models well-suited for development of novel therapeutics and diagnostics. Because of the tight linear relationship between these two parameters defining the continuous Gamma family of probability distributions, knowing one parameter enables us to infer the other, such that we can reduce the representation of data to the levels of its inherent noise-to-signal ratio to more properly use it when synthetically generating biologically inspired distributions. This approach contrasts with assuming *a priori* theoretical distributions to generate the noise portraits attempting to explain statistical variability. Our analysis offers a new standardized way to empirically capture individual and group nuances that would otherwise be discarded as gross data, thus missing important subtleties in age, genetic manipulation and treatment’s effects.

Employing empirically derived noise will be key to training new generations of biologically inspired machine learning models and artificial neural networks, to both automate classification and enable proper tracking of disease progression - an aspect of the problem that we see here in aging rats as separable from tracking progression with an intervention. This result may be relevant to both preclinical and clinical contexts. Second, we report that ABR micromovements are sensitive to human FAD-risk in both male and female rats, which could render them useful as a sensitive biomarker of AD risk very early in human life, including for tracking disease risk severity with aging. Third, we report that the standardized ABR micromovement spikes are susceptible to intervention, likely via an experience-dependent neuroplasticity mechanism. The smaller sample sizes used in this report may limit the generalizability of the findings, which require further validation in scale and in different preclinical and clinical contexts that are encouraged with the use of open source tools made available to the scientific community (Zenodo: DOI:10.5281/zenodo.17064267). Our focus on auditory neurophysiology in rats without hearing loss may raise a question about cochlear synaptopathy despite normal pure-tone thresholds that would preclude the ability to interpret effects on central auditory processing. However, it is unlikely that mutant rats had cochlear dysfunction, given the rescue of the ABR phenotype following cognitive training. Taken together, we anticipate that the translational potential of a diagnostic ABR biomarker is high, particularly since it is a rapidly acquired, pre-attentive signal collected non-invasively, which eliminates the need for prolonged or effortful subject engagement, which might be challenging in some clinical populations.

The distinct ABR signatures observed in *App^S^* rats are interpreted to reflect a general central change in neurotransmission related to AD risk that is revealed in the central auditory system. This interpretation is consistent with previous studies demonstrating that AD-related pathology disrupts synaptic transmission, neural synchrony, and temporal precision in central auditory circuits, which can manifest as an altered ABR^20,33,50–53^. Although *App^S^* rats do not exhibit classical AD histopathology at the ages studied, available data indicate the presence of functional synaptic and neurotransmission imbalances^39,41^. Importantly, that auditory learning during young adulthood normalized the ABR signature in *App^S^* rats and maintained into older age provides additional functional support for a central neural generator of initial auditory dysfunction rather than cochlear dysfunction, which would be less likely to show sustained experience-dependent recovery. Indeed, lifelong auditory learning induces experience-dependent auditory cortical neuroplasticity that can benefit neural processing and drive lasting adaptations in auditory behavior^54–56^.□Consistent with this interpretation, cortical neurons accumulate intracellular Aβ in transgenic rats expressing human APP with single (Swedish) or double mutations (e.g., Swedish and Indiana)^57–59^. Moreover, alterations in auditory evoked cortical potentials correlate with APP accumulation in the auditory cortex of APP/PS1 transgenic mice (Mo/HuAPP695swe; PS1-dE9^60^. Future studies could apply the analytical methods introduced in this report to discrete time windows of sound-evoked responses to target the role of putative brain nuclei generating the ABR along the auditory pathway that contribute to the ABR’s sensitivity to human FAD-risk.

We report in a subset of female rats after a bout of auditory-cognitive training, a transformation in *App*^S^ towards the ABR signature of a “healthy” *App*^H^ rat, despite its maintained genetic risk. The maintenance of this transformation over time and aging suggests that a central neural mechanism can reasonably promote functional resilience. An important consideration for interpreting the intervention-induced effects of auditory cognitive training is that, even though persistent, they are not expected to reverse or prevent AD-related neurodegeneration or cognitive decline. Rather, this finding provides insight into the underlying neurobiology that may be affected in parallel in both hearing function and dementia risk. Further investigation into the mechanisms underlying experience-dependent changes during intervention, and its relevance in both male and female contexts, might have promise for harnessing directly to mitigate AD risk. Notably, it is important to remember that the subjects used in these studies had no evidence of hearing loss. While there has recently been a strong representation in the field of “causal” explanations for the hearing/AD relationship, where hearing loss is assumed to directly lead to cognitive decline and/or dementia, there remains limited evidence to support or refute the link. Altogether, the findings in this report are consistent with an alternate “common cause” hypothesis for the relationship between hearing and AD/ADRD rooted in a novel functional synaptic pathology that is observable in neurophysiological recordings of evoked brain activity accessible through the auditory system.

That ABR micromovement spikes are a potential biomarker for disease risk in the *App*^S^ rat is striking because knock-in rats make no preconceived assumption about the pathogenic mechanisms nor the behavioral consequences of the knocked-in variants; only the unbiased genetic one^46^. In contrast, transgenic models, which produce, e.g., high levels of Aβ and can readily deposit amyloid plaques, are based on the hypothesis that plaques and/or other forms of toxic Aβ have a central pathogenic role in AD. Because there are known human cases where plaques can be present without dementia (e.g., in people who have other protective mutations, such as *ApoE2*), and importantly, plaques and tangles are not present in mid-life, when hearing becomes a major risk factor. Therefore, knock-in animal models may be the way of the future for hopeful lines of research to best explain the very early mid-life biological links between hearing and ADRD, independent of the amyloid cascade and/or tau pathology^61^. Consistent with the apparent functional synaptic outcome we report from the auditory system, the *App*^S^ rat has glutamate/GABA transmission imbalances^39,41^ that are now becoming more appreciated in the context of AD/ADRD with relevance to pathogenesis in cortical and subcortical structures. The relevance of ABR neurophysiology to potential cognitive phenotypes is supported by recent work in wildtype rats, showing that early ABR waves have “memory effects” that correlate with learning-induced cortical changes related to successful memory formation. The larger the change in ABR Wave amplitudes, and the larger the change in cortical tuning, and behavioral memory^1^, extending to experience-dependent plasticity related to early life stress. These ABR findings and others, including those that investigate the complex sound-evoked frequency following response (FFR)^18,64,64–66^ are consistent with memory-mediated descending cortical control on more peripheral auditory structures of the auditory system that can impose a cognitive structure on earlier neural processing in the auditory pathway to dynamically and adaptively mediate our listening abilities over a lifetime. Further, the auditory cortex is massively integrated with every subcortical processing node in the auditory system^66,67^ in descending chains of projections that outweigh the ascending inputs more than 3:1^68,69^. Thus, like other sensory systems, the auditory cortex is positioned to integrate and modulate its own ascending inputs from the periphery in a way that reflects the individual’s unique memories of salient sounds^18,70,71^. While experience-based modulation is one of its primary features, the auditory system is also a high-resolution spectrotemporal processor with the brain’s fastest, most temporally precise neural activity on the microsecond scale.

Overall, the potential for ABR micromovement spikes as a biomarker in neurodegenerative disease complements work emerging in neurodevelopmental studies that underscore the utility of the central auditory nervous system as a convenient neurophysiological window into early indications of progressive dysfunction in the brain across the lifespan^31,37^. Characteristic features of the population dynamics of sound-evoked ABRs identified in the current report may provide early, non-invasive biomarkers for detecting the severity of AD risk, and may also offer a readout to stratify AD based on, e.g., specific disease-generating mutations. It is important to note that the selection of multiparametric features that offer distinguishing characteristics between neurotypical and atypical (disease) are generated by disease-relevant population data, which enables biomarker specificity between disease states (e.g., autism vs. dementia) and within disease states (e.g., FAD mutations related to APP vs. PSEN1). The stochastic signatures of ABR micromovements are feature-specific, such that different ABR parameters (e.g., timing vs. amplitude) define distinct feature spaces with different quadrant interpretations. This rich feature space opens the door for different features to represent different features of disease state and progression or intervention. For example, leveraging the sensitivity and specificity of an enriched ABR dataset parameter space with early detection could enable timely, tailored strategies for precision medicine tailored to the individual’s underlying disease etiology and prognosis. While the current report is limited by the exclusive use of rat models of AD risk related to APP processing and small sample sizes, it provides proof-of-concept that analytical tools developed in humans effectively reverse translating into animal models across the lifespan. The report points to a new quantifiable way to uncover individualized traits, and a new way to track and interpret changes that emerge from longitudinal data. Thus, this work offers an important entry point for future investigation into other preclinical models relevant to neurodevelopmental and neurodegenerative diseases, the latter including extension into sporadic models of AD or of tauopathies in ADRD. Future studies may use ABRs to characterize functional synaptic neuropathology in models of disease related to environmental exposures or sporadic cases of AD, with or without overt hearing loss, and in longitudinal studies that can track susceptibility or resilience in parallel with cognitive phenotyping. Neurophysiological signatures that selectively distinguish dementia risk are likely to provide insights on novel etiologies to stratify AD/ADRD beyond classic clinical hallmarks for applying to next-generation human precision medicine.

## MATERIALS AND METHODS

### Experimental Design and Animal Models

#### Animal lines and genetic background

The premise of this work was to acquire electrophysiological recordings from the auditory brain of living subjects to potentially phenotype disease risk in preclinical models of human Alzheimer’s Disease (AD). Subjects were age-matched, male and female CRISPR-mediated genetic knock-in rats aged 4-5 months or 12-24 months developed to model human familial Alzheimer’s Disease (FAD) risk (breeder rats were generous gifts from Dr. Luciano D’Adamio; Rutgers New Jersey Medical School). Rats harbored the homozygous FAD Swedish risk variant that leads to early onset (<65 years old) AD in humans, here knocked into the amyloid precursor protein (*App*) rat locus, or a humanized form of *App* as non-FAD risk controls, as previously described and characterized^39–41^. Briefly, homozygous *App^S^*or *App^H^*rats carry two copies of the human Swedish risk mutation or humanized variant, respectively^39^ (N=2-4 per genotype X age), knocked into the rat *App* locus. These next-generation models make no *a priori* assumption about the pathogenic mechanisms nor the behavioral consequences of the knocked-in variants; they introduce only the unbiased genetic risk^46^. Importantly human FAD cases are often linked to autosomal dominant mutations in *App,* and genetic data suggest that APP proteolysis has a central role in both familial and sporadic AD pathogenesis^72,73^. Given that aggregated forms of Aβ are widely considered the primary pathogenic factor in AD (whether sporadic, 95% of cases, or familial, 5% of cases), we first targeted the study of an *App*^S^ model. Because human Aβ may have a higher propensity than rodent Aβ to form yet-to-be-identified toxic forms of Aβ, we contrasted the risk model with a “humanized” rat that has an Aβ component sequence (*App*^H^) complement to the *App*^S^ model. An independent cohort of knock-in 12-month-old rats (N=2 per sex) modelled an alternate FAD risk variant to the presenilin-1 (*Psen1*) locus, as previously described and characterized^42^, that—like the APP Swedish mutation—also leads to early onset AD in humans (these aged rats were generous gifts from Dr. Marc Tambini; Rutgers New Jersey Medical School). Briefly, *Psen1^LF^* rats were homozygous for the human PSEN1L435F risk variant mutation knocked into to the rat *Psen1* locusand were compared to *Psen1* wildtype non-FAD risk controls (N=1 per sex). Note that *App^H^* rats carry a wildtype *Psen1* gene and were used as non-FAD risk controls where indicated.

#### Group size

Data from a total of 17 out of 19 rats were used in the study. Sub-groupings by genotype, sex, and age produced group sizes of N=2-4 (see sample sizes per group in **SUPPLMENTARY TABLE S4**). Though the small sample sizes are a potential limitation of the generalizability of the effects reported in this study, this report is motivated by the significance of introducing feasibility and the translationally relevant potential of the analytical methods reported within.

#### Husbandry

All animals were co-housed in a colony room with a 12-hr light/dark cycle with ad libitum access to water and food. A subset of female rats were used in an intervention manipulation that implemented a behavioral training procedure (see *Behavioral Procedures*); throughout behavioral training only, rats were water-restricted with daily supplements provided to maintain an ~85% free-drinking body weight (matched to littermate controls).

#### Animal care and ethical approvals

At the termination of all studies, subjects were humanely euthanized by anesthetic overdose (Euthasol™ as 390 mg/ml sodium pentobarbital, 150mg/kg, i.p.), and death confirmed by cessation of heartbeat and decapitation. All procedures were approved and conducted in accordance with guidelines by the Institutional Animal Care and Use Committee at Rutgers, The State University of New Jersey (#PROTO999900026 to K.M.B.), in accordance with relevant named guidelines and regulations, and are reported in compliance with ARRIVE guidelines.

### Electrophysiological Recording Procedures

#### Auditory brainstem responses (ABRs)

Auditory brainstem responses (ABRs) were recorded (Medusa4Z, TDT Inc.) in anesthetized rats (ketamine(K)-xylazine(X), K: 40 mg/kg, X:4 mg/kg, i.p.) by an experimenter blind to sex and genotype using automated software (BioSigRZ, TDT Inc.) set to record single-trial trace data for analysis. Three minimally invasive subdermal electrodes (impedance 1 kΩ) were positioned at the midline along the head (recording), immediately below the left pinna (reference), and 1 cm posterior from the recording electrode (ground) in subjects whose body temperature was monitored by rectal probe and controlled to 36-37°C by a closed-loop heating blanket pad and noise-free homeothermic monitoring system (Harvard Instruments©) all located inside the sound-proof chamber. Sound stimuli were clicks and two pure-tones (5.0 kHz or 11.5 kHz; 5msec duration; 2 msec cosine-gated rise/fall time) presented via a calibrated electromagnetic speaker (MF1; Tucker Davis Technologies, Inc.; 10-90 dB SPL in 10 dB SPL steps for clicks; 20-90 dB SPL in 10 dB steps for tones) presented at a rate of 21 Hz to the right ear from a speaker positioned 4 cm away, as previously described^51,55^. Sound stimuli were presented in a blocked format (512 stimuli per block). Single-trial data used in **Data Analysis: Single-trial ABRs timeseries analysis** (described below) were evoked by 70 dB SPL, 5 msec, pure tones (512 trials per recording session) and recorded with a system cycling rate of 195,200 Hz. Neural data was bandpass filtered between 0.3 and 3.0 kHz during acquisition and stored for subsequent offline analysis using custom MATLAB© scripts. Stimulus presentation and neural response recordings were carried out using RZ6-A-P1 Multi processor, MedusaZ amplifier, and BioSig RZ software (Tucker Davis Technologies, Inc.). No additional filtering was applied offline. Click-evoked ABRs were used as an initial screen to confirm auditory function by the emergence of a characteristic trial-averaged ABR waveform in each subject online (using BioSigRZ, TDT Inc.) before acquiring tone-evoked responses (**SUPPLEMENTARY TABLE S1**). A subset of animals underwent multiple recording sessions to track training-induced effects of two-tone discrimination learning (described in **Behavioral Procedures**). In the trained cohort, auditory brainstem responses (ABRs) were recorded twice in anesthetized rats as described above to three pure-tones (5.0 kHz, 5.9 kHz and 11.5 kHz) at multiple time points: (1) Twenty-four hours prior to the first two-tone discrimination task (2TD) training session and (2) Twenty-four hours following the final 2TD training session, and (3) repeatedly up to 9 months following the final task training session. Experimenters were blind to genotype and collected data using automated software (i.e., BioSigRZ, TDT Inc.) (see experimental timeline in **SUPPLEMENTARY FIGURE S10**).

### Data Analysis

#### Trial-averaged ABRs

The trial-averaged ABR evoked by clicks or pure tones (5kHz or 11.5 kHz, 10-90 dB SPL in 10 dB SPL steps for clicks; 20-90 dB SPL in 10 dB steps for tones) amplitude were used to estimate each subject’s hearing threshold (in dB SPL) (**SUPPLEMENTARY TABLE S1**), and to analyze the tone-evoked (5kHz, 70 dB SPL) response in the second positive wave of the ABR (PW2), which appears as a positive peak in the trial-averaged waveform, and is reliably observed in both naïve, aged, and trained rats^62,74^ (**SUPPLEMENTARY FIGURE S11; SUPPLEMENTARY TABLES S2, S3)**. Hearing thresholds were determined by concordance between two independent and experienced human raters, defined as the lowest sound level (in dB SPL) to elicit a characteristic ABR waveform in the tone-evoked trial-averaged response (**SUPPLEMENTARY FIGURES S1, S2**). Hearing thresholds were defined from trial-averaged ABR traces as the lowest sound level eliciting a visually identifiable ABR waveform identified by a visually-detectable wave 2 (PW2; see **SUPPLEMENTARY FIGURES S2, S11**) as that characteristic tended to be the most reliable peak across the recordings and in studies of intervention-induced effects used previously to detect experiential effects^62,74^. Thresholds were confirmed by inter-rater reliability between at least two independent human raters. In cases of non-concordance, a third independent and experienced human rater made an assessment to ensure at least 2/3 rater concordance for final hearing threshold determination in all subjects. All raters were blind to subject sex, genotype, and age. The resulting ABR thresholds (in dB SPL) were compared using unpaired t-tests with Welch’s correction (α = 0.05) for between-group analyses (genotype, sex, and age) and paired t-tests (α = 0.05) for within-group comparisons (trained vs. untrained) *App^S^* and *App^H^ rats.* Custom Matlab scripts were used to identify peaks within the trial-averaged tone-evoked ABR waveform and to derive the trough-to-peak wave amplitude (uV). Wave amplitude was defined as the voltage difference between the preceding trough and the subsequent peak in the trial-averaged ABR (**SUPPLEMENTARY FIGURE S11)**. **SUPPLEMENTARY TABLE S3** shows conventional ABR metrics of amplitude and latency for tone-evoked responses. Further, two-tailed Pearson correlation analyses were used to assess relationships between experiential-related changes in the PW2 amplitude and the success of task memory formation (see *Brain-Behavior correlational analysis*, below). All statistical analyses on trial-averaged ABRs were performed using GraphPad Prism software (Boston, Massachusetts, USA).

#### Single-trial ABRs timeseries analysis

Populations of single-trial ABRs were analyzed using mathematical modeling of the distributions of single trial electrophysiological trace features across 100s of repeated sound presentations, the mathematics of which have been described in detail^31,34^ and scripts have been made available for *any* continuous time-series analysis on Zenodo (<u>record 6299560#.Y3las3bMLcs</u>)^31^. A continuous timeseries of trial-by-trial micromovements, which are small fluctuations in neural activations within each ABR trial away from the empirically derived mean and normalized to scale out allometric effects of anatomically disparate sizes across animals, was used to construct a multidimensional parameter space for analysis. Such individualized trial fluctuations away from the non-stationary empirical mean, are typically discarded in conventional trial-averaging approaches that assume a theoretical distribution. Here such nuances in the trial-by-trial data were retained to capture rich features of signal variability. Briefly, this method was originally developed with inspiration from the timeseries motor movement datasets across repeated trials of voluntarily reaching (e.g., for objects)^37^. They were later generalized across a multiplicity of biorhythms of the nervous systems (eyes, face, voice, heart, voice, neural signals)^75^ and dynamic interrogation of human transcriptome data^76^. By eye, these signals appear smooth and continuous. They however have moment-to-moment variations over the shifting empirical mean, that over time inform the system of emerging patterns ranging from random and unpredictable, to organized and highly predictable, such that past events inform of future events. As in a moving average, it is possible to empirically determine the mean peaks and valleys in the signals of interest (as opposed to taking a theoretical mean assumed from an *a priori* imposed distribution). Then, as the system senses the stimuli and responds, the response fluctuations relative to the empirically estimated mean can be tracked moment-by-moment. A standardization of these fluctuations away from the mean to account for possible allometric effects due to anatomical disparities (such as differences in head circumference and end-effectors’ lengths across animals), produce the so-called “micromovement spikes” with micro-peaks. Continuous time series data across trials generates thousands of such micro-peaks whose distribution parameters can be empirically estimated using e.g., maximum likelihood estimation. As an example, plotting the empirical parameters thus estimated with 95% confidence intervals on a parameter space automatically separated the auditory brainstem responses (ABR) in human neonates distinguishing even 5dB differences in audible and non-audible stimuli. The responses separated those babies who went on to develop neurotypically from those with neuro-atypical development receiving a diagnosis of autism spectrum disorders 4 years later ^31^. Here, we adapt the methods to neurophysiological ABR data from a rat model of disease risk. It should be understood then that such normalized micro-peaks reflect small fluctuations in neural activations and at a value of 0 represent local mean activity (**FIGURE 1**). Across biophysical signals, the continuous Gamma family of probability distribution functions has proved to be the best fit (in an MLE sense), capturing human maturational^37^ and neurodegenerative^77^ stages in the form of a scaling power law. ABR “micromovements spikes” reflect characteristics of synaptic transmission along the central auditory pathways, which are usually unobservable in standard trial-averaging practices. Importantly, these patterns are specific to each animal but because of the normalization for allometric effects, the distribution parameters of standardized micro-peaks can be compared on the same parameter space across multiple animals, e.g., within or between groups. In this way it is not only possible to quantify individual shifts in probability distributions but also to capture automatically self-emerging classes of animals with similar stochastic properties. Similarity here is ascertained with a proper distance metric operating on points of a probability distribution function space. Since each animal spans a whole family of PDFs, this method captures what turned out to be important nuanced information, otherwise thrown away as gross data.

**Equation 1** provides the proper normalization of the peaks that deviate from the empirically estimated mean:

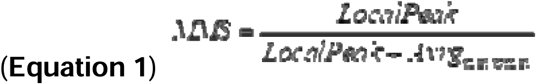

This approach enabled optimal feature extraction and the derivation of interpretable parameter spaces for downstream comparisons between individuals and experimental groups. Effects of aging, genotyping and treatment were possible to ascertain using this new statistical platform for individualized response analysis.

### Behavioral Procedures

All behavioral training and memory test sessions were conducted in instrumental conditioning chambers within a sound-attenuated box (Habitest Modular System, Coulbourn Instruments©). Rats were water-restricted, handled and initially habituated to the training procedures by learning to bar press for water reward (0.1cc) in five 45-60 minute bar-press shaping sessions before beginning Auditory Training as shown in the experimental timeline (**SUPPLEMENTARY FIGURE S5**). This Handling/Habituation phase of training ensured that all animals were trainable and could safely acquire the procedural aspects of the task (i.e., bar-pressing for rewards) before any sounds were introduced. Next, rats were trained to bar press only to pure tones for reward in a *two-tone* discrimination task (2TD). Responses to a target pure tone frequency (S+, 5.0 kHz; 75±5 dB SPL, 8 s) resulted in reward, while responses to the non-target (S-, 11.5 kHz; 75±5 dB SPL, 8 s) were unrewarded and triggered an error signal (flashing house light) and a “time-out” (an extended 6 s interval until the start of the next trial). It is important to note that the 2TD task is not difficult perceptually since the acoustic frequencies (5.0 kHz vs. 11.5 kHz) are over an octave apart and easily distinguished by naïve rodents^78^. Rather, the behavioral challenge was associative in nature: 2TD performance demands the formation of memory for which tone frequency is associated with reward (vs. no reward). The 5.0 kHz pure tone was assigned as the Go (rewarded) stimulus, while 11.5 kHz served as the No-go stimulus without counterbalancing to avoid any confounds of frequency-specific effects. As the data were not normally distributed, Mann–Whitney U test was used to verify that the 11.5 kHz tone-evoked ABR thresholds were stable between recordings made before vs. after training. There was no significant in hearing thresholds, including for the higher frequency No-go 11.5 kHz tone (Mann–Whitney U test, p > 0.05; **SUPPLEMENTARY TABLE S1**). Inter-trial intervals (ITIs) were an average of 8 s (range: 4–12 s intervals, randomized across trials). Bar presses during an ITI were inconsequential (i.e., no time-out, no reward). 2TD training sessions were 30-40 minutes in length in daily sessions Monday through Friday. Subjects had Saturday and Sunday without training with ad libitum access to food and water that ended midday on Sundays. At that time and throughout training, water restriction was in place and water supplementation administered to maintain ~85% free-drinking body weight throughout all behavioral training and memory testing. If needed, water supplementation occurred no sooner than 2 hours after a single training session to avoid motivational effects of free drinking water during task performance.

#### Behavioral data analysis

Daily *two-tone* discrimination (2TD) performance was calculated as before^79^ as the number of bar presses (BP) to the rewarded tone (S□+)□divided by the number of bar presses to both the S+ and unrewarded tone (S−) as follows: P□=□100% x [#BP^S+^/ (BP^S+^ + #BP^S−^)]. 2TD training continued until rats reached performance asymptote or to a maximum of 20 training sessions. The performance asymptote criterion was defined by the combination of: (1) performance ≥70%, (2) ≤10% coefficient of variation across 3 consecutive days, and (3) [(# BP^S+^/ #Trials^S+^) - (# BP^S−^/# Trials^S−^)] / [(# BP^S+^/# Trials^S+^) + (# BP^S−^ / # Trials^S−^)] ≥1. Twenty-four hours following the final asymptotic 2TD training session, rats were given a tone memory test to determine the success of auditory memory formation after task training. During the memory test, rats were presented with individual trials of sound presentations across different acoustic frequencies, including the S+ and S− tones. Ten different pure tone frequencies were tested: 3.6, 4.2, 5.0 (S+ rewarded tone), 5.9, 7.0, 8.3, 9.7, 11.5 (S− unrewarded tone), 13.6, and 16.0□kHz (all pure tones presented at a sound level of 75±5 dB SPL). The memory test began with 15 regular trials of the 2TD task (i.e., bar presses to the S+ tone frequency were rewarded; bar presses to the S− were not) to confirm stable task performance on the day of the memory test. Each tone frequency was presented a total of 8 times. The session was divided into two continuous blocks, with two presentations of each tone per block in a pseudorandom order. No responses were reinforced. All animals were confirmed to show a response profile across the 10 test tone frequencies consistent with 2TD training as a tendency to respond more to lower acoustic frequencies around the S+ and less to higher frequencies around the S-, which confirmed general hearing and learning ability. To quantify the success of tone-reward memory formation for the specific frequency (to within ¼ octave of the rewarded signal tone), a tone memory index was calculated as the proportion of responses made to the S+ tone (5.0 kHz) during the memory test, minus the proportion of responses made to the immediately neighboring test tone frequency (5.9 kHz). The more positive the value, the more successful the memory for rewarded S+ tone frequency (vs. a nearby neighbor frequency) for that subject^74^.

#### Brain-behavior correlational analysis

Contrast measures were derived to quantify the specificity of intervention-induced differences in the PW2 amplitude evoked by 5.0 kHz (the trained S+ tone, 70 dB SPL) vs. by 5.9 kHz (70 dB SPL), a novel nearby (within ¼ octave) neighboring tone frequency, after learning the 2TD task. A contrast measure was calculated within each subject as the difference between frequencies in post-minus pre-training PW2 amplitudes. As such, a more positive value indicates a relatively larger increase in the PW2 amplitude evoked by the S+ tone (5.0 kHz). Pearson correlation analysis was used to evaluate the relationship between intervention-induced increases in PW2 amplitude evoked by 5.0 kHz (vs. 5.9 kHz) (**SUPPLEMENTARY FIGURE S11C**) and the preference to behaviorally respond to the S+ tone (5.0 kHz) during the memory test, which was previously reported to track with the success of memory formation and related auditory cortical plasticity^62,74^.

### Inclusion and Exclusion Criteria

All rats underwent a screening phase for inclusion in the single-trial ABR timeseries datasets to ensure the reliability of single-trial data acquired across the entire duration of the recording session for accurate within-animal phenotyping and generalizability of the results across the group. To do so, we divided continuous time series data within each individual subject into 8 equal epochs and estimated the trajectory of data distribution and confidence intervals of the measured parameters within the subject’s recording session. A Noise-to-Signal Ratio (NSR) was then calculated by dividing the variance of the electrophysiological signal by its mean. This approach allowed us to determine within-animal trajectories reflecting the consistency of data acquired within the same recording session, to assess changes in distributional characteristics, and compare these patterns across rats within the same experimental group (e.g., genotype). One rat (female *App^S^*) exhibited a markedly different NSR trend from the rest of the group by showing an orthogonal trajectory and was a clear outlier of the group profile. To prevent a reduction in the generalizability of our results, this rat was excluded from analysis. An additional rat (female *App^H^*) showed aberrant behavior, so was also excluded from analysis. In total, data from 17 of 19 rats studied were included. All screening procedures were conducted blind to genotype and sex. All animals and data included in the intervention study to determine effects of auditory training passed the inclusion criteria.

## Acknowledgments

We thank Dr. Marc Tambini and Dr. Luciano D’Adamio for their generous gifts of experimental animals and breeder knock-in rats, which supported the establishment of our colony and the completion of this work. We also thank all members of the CLEF Lab, especially Nilay Ateşyakar, at Rutgers University and Dr. Michal Beeri at the Herbert and Jacqueline Krieger Klein Alzheimer’s Research Center for their continuous insight and valuable input into this project.

## Author Contributions

Conceptualization: A.G.A. and K.M.B.; Methodology: A.G.A. and K.M.B.; Investigation: A.G.A.; E.Y., P.M., F.R.; Visualization: A.G.A., E.B.T. and K.M.B.; Data analysis: A.G.A., P.M., F.R., E.Y., S.R., E.B.T. and K.M.B.; Funding acquisition: K.M.B.; Supervision: K.M.B., (computational) E.B.T.; Software: E.B.T.; Writing – original draft: K.M.B. and A.G.A.; Writing—review and editing: A.G.A., E.B.T., and K.M.B. All authors read and approved the manuscript.

## Data Availability Statement

Data are available in a Zenodo data repository (https://zenodo.org/records/17064267; DOI:10.5281/zenodo.17064267). The code used to generate analytical methods in rodents was adopted from code originally developed for human data available in Zenodo record (https://zenodo.org/records/6299560#.Y3las3bMLcs).

## Funding

This work was supported by the National Institutes of Health, the National Institute on Deafness and Communication Disorders [R01-DC018561 to K.M.B.]. E.B.T. was funded by the Nancy Lurie Marks Family Foundation Career Continuation Award.

## Competing interests

The authors declare no competing interests.

## Notes

### Competing Interest Statement

The authors have declared no competing interest.

### Summary of Updates

Multiple sections throughout the manuscript have been revised for clarity; small corrections and additions have been made to each of the four main figured; additional analyses have been included to complement and provide further support for the original sets of interpretation; and figures added to the supplemental material. The Discussion has been lengthened to better address limitations of the study.

## REFERENCES

1. Panza, F. et al. Age-related hearing impairment and frailty in Alzheimer’s disease: interconnected associations and mechanisms. Front. Aging Neurosci. 7, (2015).

2. Lin, F. R. et al. Hearing Loss and Incident Dementia. Arch. Neurol. 68, (2011).

3. Livingston, G. et al. Dementia prevention, intervention, and care. The Lancet 390, 2673–2734 (2017).

4. Taljaard, D. S., Olaithe, M., Brennan-Jones, C. G., Eikelboom, R. H. & Bucks, R. S. The relationship between hearing impairment and cognitive function: a meta-analysis in adults. Clin. Otolaryngol. 41, 718–729 (2016).

5. Mamo, S. K. et al. Hearing Care Intervention for Persons with Dementia: A Pilot Study. Am. J. Geriatr. Psychiatry 25, 91–101 (2017).

6. Maharani, A. et al. Longitudinal Relationship Between Hearing Aid Use and Cognitive Function in Older Americans. J. Am. Geriatr. Soc. 66, 1130–1136 (2018).

7. Dawes, P. et al. Hearing Loss and Cognition: The Role of Hearing Aids, Social Isolation and Depression. PLOS ONE 10, e0119616 (2015).

8. Lin, F. R. & Albert, M. Hearing Loss and Dementia – Who’s Listening? Aging Ment. Health 18, 671–673 (2014).

9. Lin, F. R. et al. Hearing intervention versus health education control to reduce cognitive decline in older adults with hearing loss in the USA (ACHIEVE): a multicentre, randomised controlled trial. Lancet Lond. Engl. 402, 786–797 (2023).

10. Reed, N. S. et al. Recruitment and baseline data of the Aging and Cognitive Health Evaluation in Elders (ACHIEVE) study: A randomized trial of a hearing loss intervention for reducing cognitive decline. Alzheimers Dement. N. Y. N 10, e12453 (2024).

11. Weinberger, N. M. & Bieszczad, K. M. Introduction: From Traditional Fixed Cortical Sensationism to Contemporary Plasticity of Primary Sensory Cortical Representations. in Neurobiology of Sensation and Reward (ed. Gottfried, J. A.) (CRC Press/Taylor & Francis, Boca Raton (FL), 2011).

12. Bieszczad, K. M. & Weinberger, N. M. Representational gain in cortical area underlies increase of memory strength. Proc. Natl. Acad. Sci. U. S. A. 107, 3793–3798 (2010).

13. Bieszczad, K. M. & Weinberger, N. M. Extinction reveals that primary sensory cortex predicts reinforcement outcome. Eur. J. Neurosci. 35, 598–613 (2012).

14. Schreiner, C. E. & Polley, D. B. Auditory map plasticity: diversity in causes and consequences. Curr. Opin. Neurobiol. 24, 143–156 (2014).

15. Pienkowski, M. & Eggermont, J. J. Cortical tonotopic map plasticity and behavior. Neurosci. Biobehav. Rev. 35, 2117–2128 (2011).

16. McGann, J. P. Associative learning and sensory neuroplasticity: how does it happen and what is it good for? Learn. Mem. Cold Spring Harb. N 22, 567–576 (2015).

17. Noda, T., Aschauer, D. F., Chambers, A. R., Seiler, J. P.-H. & Rumpel, S. Representational maps in the brain: concepts, approaches, and applications. Front. Cell. Neurosci. 18, 1366200 (2024).

18. Kraus, N. & White-Schwoch, T. Unraveling the biology of auditory learning: A cognitive-sensorimotor-reward framework. Trends Cogn. Sci. 19, 642–654 (2015).

19. Lesicko, A. M. H. & Geffen, M. N. Diverse functions of the auditory cortico-collicular pathway. Hear. Res. 425, 108488 (2022).

20. Gates, G. A., Anderson, M. L., Feeney, M. P., McCurry, S. M. & Larson, E. B. Central Auditory Dysfunction in Older People with Memory Impairment or Alzheimer’s Dementia. Arch. Otolaryngol. Head Neck Surg. 134, 771–777 (2008).

21. Johnson, J. C. S. et al. Hearing and dementia: from ears to brain. Brain J. Neurol. 144, 391–401 (2021).

22. Parkkonen, L., Fujiki, N. & Mäkelä, J. P. Sources of auditory brainstem responses revisited: Contribution by magnetoencephalography. Hum. Brain Mapp. 30, 1772–1782 (2009).

23. Chandrasekaran, B. & Kraus, N. The scalp-recorded brainstem response to speech: neural origins and plasticity. Psychophysiology 47, 236–246 (2010).

24. Oyler, R. F., Lauter, J. L. & Matkin, N. D. Intrasubject variability in the absolute latency of the auditory brainstem response. J. Am. Acad. Audiol. 2, 206–213 (1991).

25. Bidelman, G. M., Pousson, M., Dugas, C. & Fehrenbach, A. Test-Retest Reliability of Dual-Recorded Brainstem versus Cortical Auditory-Evoked Potentials to Speech. J. Am. Acad. Audiol. 29, 164–174 (2018).

26. Auditory brainstem response. in Handbook of Clinical Neurology vol. 160 451–464 (Elsevier, 2019).

27. Santos, M. A. R., Munhoz, M. S. L., Peixoto, M. A. L. & Silva, C. S. High click stimulus repetition rate in the auditory evoked potentials in multiple sclerosis patients with normal MRI. Does it improve diagnosis. Rev. Laryngol. - Otol. - Rhinol. 125, 151–155 (2004).

28. Xing, Y. et al. Age-related changes of myelin basic protein in mouse and human auditory nerve. PloS One 7, e34500 (2012).

29. Kraus, N. et al. Subconcussion revealed by sound processing in the brain. Exerc. Sport Mov. 1, 1–4 (2023).

30. Skoe, E. & Chandrasekaran, B. The layering of auditory experiences in driving experience-dependent subcortical plasticity. Hear. Res. 311, 36–48 (2014).

31. Torres, E. B. et al. Sensing echoes: temporal misalignment in auditory brainstem responses as the earliest marker of neurodevelopmental derailment. PNAS Nexus 2, 315 (2023).

32. Patel, S. P. et al. Neural Processing of Speech Sounds in ASD and First-Degree Relatives. J. Autism Dev. Disord. 53, 3257–3271 (2023).

33. Tarawneh, H. Y., Mulders, W. H. A. M., Sohrabi, H. R., Martins, R. N. & Jayakody, D. M. P. Investigating Auditory Electrophysiological Measures of Participants with Mild Cognitive Impairment and Alzheimer’s Disease: A Systematic Review and Meta-Analysis of Event-Related Potential Studies. J. Alzheimers Dis. JAD 84, 419–448 (2021).

34. Vander Werff, K. R. & Rieger, B. Brainstem Evoked Potential Indices of Subcortical Auditory Processing After Mild Traumatic Brain Injury. Ear Hear. 38, e200–e214 (2017).

35. Irimajiri, R., Golob, E. J. & Starr, A. Auditory brain-stem, middle- and long-latency evoked potentials in mild cognitive impairment. Clin. Neurophysiol. Off. J. Int. Fed. Clin. Neurophysiol. 116, 1918–1929 (2005).

36. Wilson, W. J., Penn, C., Saffer, D. & Aghdasi, F. Improving the prediction of outcome in severe acute closed head injury by using discriminant function analysis of normal auditory brainstem response latencies and amplitudes. J. Neurosurg. 97, 1062–1069 (2002).

37. Torres, E. B. et al. Autism: the micro-movement perspective. Front. Integr. Neurosci. 7, 32 (2013).

38. Vidler, M. & Parkert, D. Auditory brainstem response threshold estimation: subjective threshold estimation by experienced clinicians in a computer simulation of the clinical test. Int. J. Audiol. 43, 417–429 (2004).

39. Tambini, M. D., Yao, W. & D’Adamio, L. Facilitation of glutamate, but not GABA, release in Familial Alzheimer’s APP mutant Knock-in rats with increased β-cleavage of APP. Aging Cell 18, e13033 (2019).

40. Tambini, M. D., Norris, K. A. & D’Adamio, L. Opposite changes in APP processing and human Aβ levels in rats carrying either a protective or a pathogenic APP mutation. eLife 9, e52612.

41. Yesiltepe, M. et al. Late-long-term potentiation magnitude, but not Aβ levels and amyloid pathology, is associated with behavioral performance in a rat knock-in model of Alzheimer disease. Front. Aging Neurosci. 14, 1040576 (2022).

42. Tambini, M. D. & D’Adamio, L. Knock-in rats with homozygous PSEN1L435F Alzheimer mutation are viable and show selective γ-secretase activity loss causing low Aβ40/42 and high Aβ43. J. Biol. Chem. 295, 7442–7451 (2020).

43. Hanes, J. et al. Rat tau proteome consists of six tau isoforms: implication for animal models of human tauopathies. J. Neurochem. 108, 1167–1176 (2009).

44. Xia, D. et al. Presenilin-1 knockin mice reveal loss-of-function mechanism for familial Alzheimer’s disease. Neuron 85, 967–981 (2015).

45. Tambini, M. D. et al. Aβ43 levels determine the onset of pathological amyloid deposition. J. Biol. Chem. 299, 104868 (2023).

46. D’Adamio, L. Transfixed by transgenics: how pathology assumptions are slowing progress in Alzheimer’s disease and related dementia research. EMBO Mol. Med. 15, e18479 (2023).

47. Livingston, G. et al. Dementia prevention, intervention, and care: 2024 report of the Lancet standing Commission. The Lancet 404, 572–628 (2024).

48. van der Kant, R., Goldstein, L. S. B. & Ossenkoppele, R. Amyloid-β-independent regulators of tau pathology in Alzheimer disease. Nat. Rev. Neurosci. 21, 21–35 (2020).

49. Frontiers | Dynamic Interrogation of Stochastic Transcriptome Trajectories Using Disease Associated Genes Reveals Distinct Origins of Neurological and Psychiatric Disorders. https://www.frontiersin.org/journals/neuroscience/articles/10.3389/fnins.2022.884707/full.

50. Uhlhaas, P. J. & Singer, W. Neural Synchrony in Brain Disorders: Relevance for Cognitive Dysfunctions and Pathophysiology. Neuron 52, 155–168 (2006).

51. Gurevicius, K., Lipponen, A. & Tanila, H. Increased cortical and thalamic excitability in freely moving APPswe/PS1dE9 mice modeling epileptic activity associated with Alzheimer’s disease. Cereb. Cortex 23, 1148–1158 (2013).

52. Liu, Y. et al. Hearing loss is an early biomarker in APP/PS1 Alzheimer’s disease mice. Neurosci. Lett. 717, 134705 (2020).

53. Hamza, Y. et al. Auditory brainstem responses as a biomarker for cognition. *Commun*. Biol. 7, 1653 (2024).

54. Schreiner, C. E. & Winer, J. A. Auditory cortex mapmaking: principles, projections, and plasticity. Neuron 56, 356–365 (2007).

55. Weinberger, N. M. New perspectives on the auditory cortex: learning and memory. Handb. Clin. Neurol. 129, 117–147 (2015).

56. Aschauer, D. & Rumpel, S. The Sensory Neocortex and Associative Memory. Curr. Top. Behav. Neurosci. 37, 177–211 (2018).

57. Echeverria, V. et al. Rat transgenic models with a phenotype of intracellular Abeta accumulation in hippocampus and cortex. J. Alzheimers Dis. JAD 6, 209–219 (2004).

58. Lopez, E. M., Bell, K. F. S., Ribeiro-da-Silva, A. & Cuello, A. C. Early changes in neurons of the hippocampus and neocortex in transgenic rats expressing intracellular human a-beta. J. Alzheimers Dis. JAD 6, 421–431; discussion 443-449 (2004).

59. Folkesson, R. et al. A transgenic rat expressing human APP with the Swedish Alzheimer’s disease mutation. Biochem. Biophys. Res. Commun. 358, 777–782 (2007).

60. Mei, L., Liu, L.-M., Chen, K. & Zhao, H.-B. Early Functional and Cognitive Declines Measured by Auditory-Evoked Cortical Potentials in Mice With Alzheimer’s Disease. Front. Aging Neurosci. 13, (2021).

61. Yesiltepe, M., Yin, T. & D’Adamio, L. Beyond amyloid: altered gene function in neurodegenerative diseases. Aging 15, 9235–9237 (2023).

62. Rotondo, E. K. & Bieszczad, K. M. Precise memory for pure tones is predicted by measures of learning-induced sensory system neurophysiological plasticity at cortical and subcortical levels. Learn. Mem. Cold Spring Harb. N 27, 328–339 (2020).

63. Rotondo, E. K. & Bieszczad, K. M. Memory Specific to Temporal Features of Sound Is Formed by Cue-Selective Enhancements in Temporal Coding Enabled by Inhibition of an Epigenetic Regulator. J. Neurosci. Off. J. Soc. Neurosci. 41, 9192–9209 (2021).

64. Kraus, N., Strait, D. & Parbery-Clark, A. Cognitive factors shape brain networks for auditory skills: spotlight on auditory working memory. Ann. N. Y. Acad. Sci. 1252, 100–107 (2012).

65. Coffey, E. B. J. et al. Evolving perspectives on the sources of the frequency-following response. Nat. Commun. 10, 5036 (2019).

66. Chandrasekaran, B., Skoe, E. & Kraus, N. An integrative model of subcortical auditory plasticity. Brain Topogr. 27, 539–552 (2014).

67. Coomes Peterson, D. & Schofield, B. R. Projections from auditory cortex contact ascending pathways that originate in the superior olive and inferior colliculus. Hear. Res. 232, 67–77 (2007).

68. Asilador, A. & Llano, D. A. Top-Down Inference in the Auditory System: Potential Roles for Corticofugal Projections. Front. Neural Circuits 14, 615259 (2020).

69. Antunes, F. M. & Malmierca, M. S. Corticothalamic Pathways in Auditory Processing: Recent Advances and Insights From Other Sensory Systems. Front. Neural Circuits 15, 721186 (2021).

70. Fritz, J., Elhilali, M. & Shamma, S. Active listening: task-dependent plasticity of spectrotemporal receptive fields in primary auditory cortex. Hear. Res. 206, 159–176 (2005).

71. Fu, V. X. et al. Perception of auditory stimuli during general anesthesia and its effects on patient outcomes: a systematic review and meta-analysis. Can. J. Anaesth. J. Can. Anesth. 68, 1231–1253 (2021).

72. Bayer, T. A., Cappai, R., Masters, C. L., Beyreuther, K. & Multhaup, G. It all sticks together--the APP-related family of proteins and Alzheimer’s disease. Mol. Psychiatry 4, 524–528 (1999).

73. van der Kant, R. & Goldstein, L. S. B. Cellular functions of the amyloid precursor protein from development to dementia. Dev. Cell 32, 502–515 (2015).

74. Rotondo, E. K. & Bieszczad, K. M. Sensory cortical and subcortical auditory neurophysiological changes predict cue-specific extinction behavior enabled by the pharmacological inhibition of an epigenetic regulator during memory formation. Brain Res. Bull. 169, 167–183 (2021).

75. Torres, E. B., Cole, J. & Poizner, H. Motor output variability, deafferentation, and putative deficits in kinesthetic reafference in Parkinson’s disease. Front. Hum. Neurosci. 8, 823 (2014).

76. Bermperidis, T., Schafer, S., Gage, F. H., Sejnowski, T. & Torres, E. B. Dynamic Interrogation of Stochastic Transcriptome Trajectories Using Disease Associated Genes Reveals Distinct Origins of Neurological and Psychiatric Disorders. Front. Neurosci. 16, 884707 (2022).

77. Torres, E. B., Caballero, C. & Mistry, S. Aging with Autism Departs Greatly from Typical Aging. Sensors 20, 572 (2020).

78. Talwar, S. K. & Gerstein, G. L. Auditory frequency discrimination in the white rat. Hear. Res. 126, 135–150 (1998).

79. Shang, A., Bylipudi, S. & Bieszczad, K. M. Inhibition of histone deacetylase 3 via RGFP966 facilitates cortical plasticity underlying unusually accurate auditory associative cue memory for excitatory and inhibitory cue-reward associations. Behav. Brain Res. 356, 453–469 (2019).

