## Supplemental Materials for "Sound-evoked auditory neurophysiological signals are a window into prodromal functional differences in a preclinical model of Alzheimer’s Disease"

### Authors and Affiliations

1. **Biomedical Research Institute of New Jersey, Cedar Knolls, NJ, 07927, USA**
   Aysegul Gungor Aydin
2. **Atlantic Health System, Morristown, NJ, 07960, USA**
   Aysegul Gungor Aydin
3. **Rutgers Robert Wood Johnson Medical School, New Brunswick, NJ, 08901, USA**
   Pranav Manoj
4. **Department of Psychology—Behavioral and Systems Neuroscience, Rutgers University, Piscataway, NJ, 08854, USA**
   Elias Youssef, Sarah Rajan, Faiza Ramadan & Kasia M. Bieszczad
5. **Department of Psychology—Cognitive Psychology, Rutgers University, Piscataway, NJ, 08854, USA**
   Elizabeth B Torres
6. **Computational Biomedicine Imaging and Modelling, Rutgers University, Piscataway, Piscataway, NJ, 08854, USA**
   Elizabeth B Torres
7. **Rutgers Center for Cognitive Science (RuCCS), Rutgers University**
   Kasia M. Bieszczad and Elizabeth B Torres
8. **Department of Head & Neck Surgery and Communication Sciences, Rutgers Robert Wood Johnson Medical School, New Brunswick, NJ, 08901, USA**
   Kasia M. Bieszczad

### Corresponding author

Correspondence to Dr. Kasia M. Bieszczad
327 Psychology Building, Rutgers University-New Brunswick
152 Frelinghuysen Road
Piscataway, New Jersey 08854 USA


### SUPPLEMENTARY METHODS

**Pipeline to derive the micro-movement spikes and Gamma representation from positively rectified ABR signals.** Raw auditory brainstem response (ABR) data traces from single trials of sound evoked responses are transformed in multiple steps across an analytical pipeline (for a visual representation of the data transformations that occur in the analytical processing of single-trial ABR data, see **FIGURE 1A** in main text). The original data (**Step 1**) is used to obtain sliding windows of 1 second with 50% overlap. The peaks of the original time series in each window are then gathered into a frequency histogram and the empirical estimation of the best continuous family of probability distributions obtained. This is done according to maximum likelihood estimation using different continuous families of distributions and settling on the optimal one, with 95% confidence. The first moment (the mean is retained). This empirical estimation across windows of data gives the continuous Gamma family of probability distribution functions, which has 2 parameters, the shape and the scale. We then take the first moment of the distribution thus obtained (the empirical mean) (black line in **Step 2**) and measure the absolute deviations of each point in the original time series from the empirically estimated mean. This is a new time series depicted in **Step 3** that retains the original time stamps but now focuses on the deviations from the mean. These absolute deviations are then passed through Equation (1) above in **Step 4**, to scale out possible allometric effects due to anatomical disparities across animals. The resulting waveform in **Step 5** is the micromovement spikes. They are called micromovements because the values of the time series move away from the empirical mean and spikes is because we can study the continuous range in the real-number range [0,1] of deviations from the empirical mean, or we can study the binary data from each normalized peak as a continuous Gamma or as a Poisson process. Importantly, 0 values are at mean level, such that our analysis focuses on departures from mean activity – not on instrumentation noise. The next step (**FIGURE 1B**) provides a representation of the data in the Gamma parameter plane spanned by the shape and scale (dispersion) Gamma parameters. Here two representative rat subjects are shown as points on this plane with 95% confidence intervals for each parameter dimension. Their ABR data went through *Steps 1-5* together and show relative physiological dissimilarity in state space. 95% confidence intervals are shown using target marks around mean values.

### SUPPLEMENTARY DATA

All data within the manuscript are publicly available on Zenodo: 10.5281/zenodo.17064267.

### SUPPLEMENTARY FIGURES

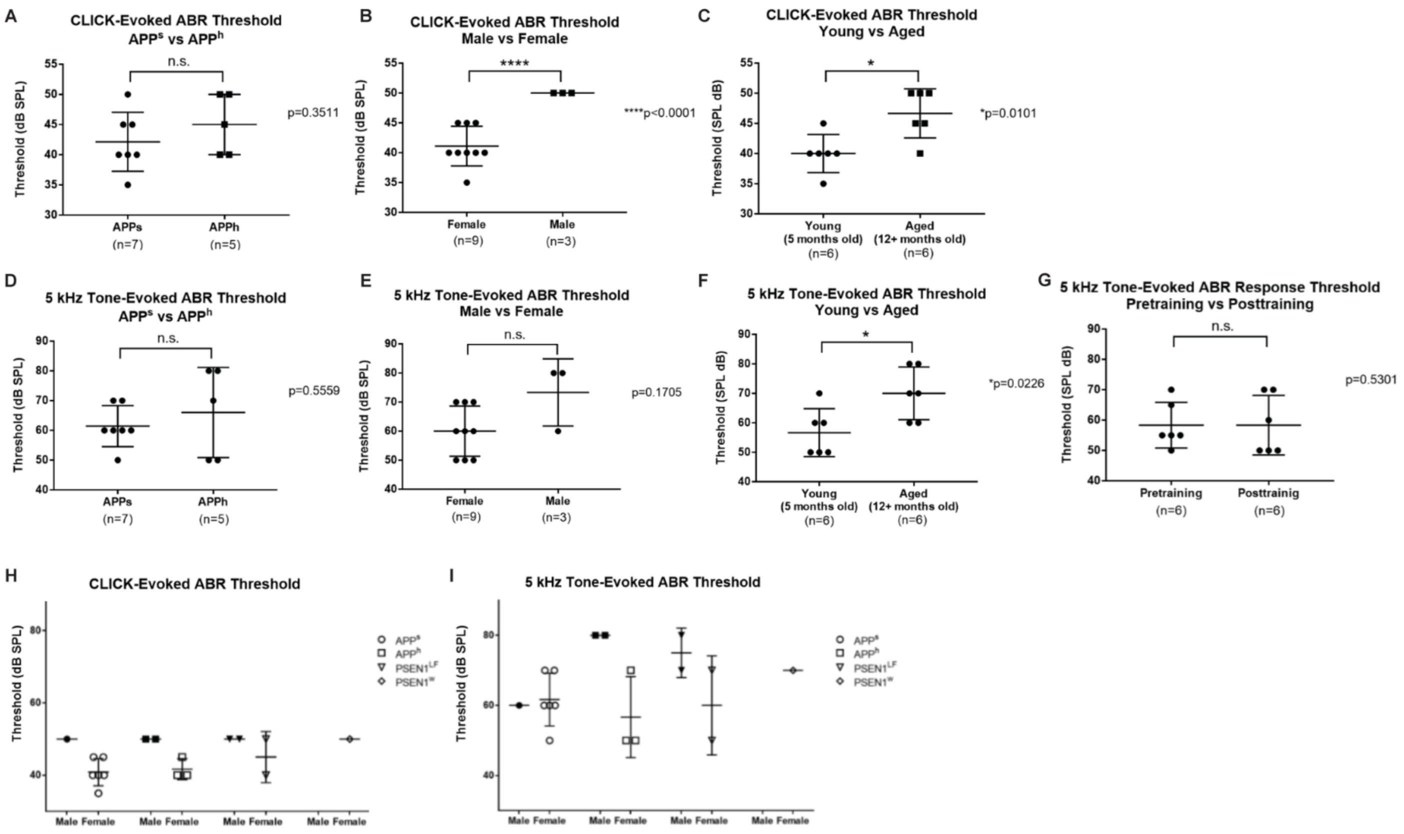


**Supplemental Figure S1:** *Tone and click evoked Auditory brainstem response (ABR) hearing thresholds across genotype, sex, age, and training*. ABR thresholds (for both tone and click stimuli) were determined from all animals included in the study (N=17). The cohort was subsequently divided only for specific analyses (genotype, age, and sex comparisons) for between-group comparisons. **(A–C)** Click-evoked ABR thresholds comparing *APP* genotype ($App^{s}$ and $App^{H}$ rats), sex (female vs. male), and age (young: 5 months; aged: 12+ months). **(D–F)** 5 kHz tone-evoked ABR thresholds for the same comparisons in A to C, as well as **(G)** before and after sound-reward training. **(H)** Individual click-evoked and 5 kHz tone-evoked **(I)** ABR thresholds plotted by genotype and sex (symbols indicate genotypes: $App^{s}$, $App^{H}$, $PSEN1^{LF}$, $PSEN1^{w}$). Each symbol represents individual animal; horizontal lines indicate group mean ± SEM. Sample sizes are shown beneath each group. Statistical significance is indicated above comparisons (*$p < 0.05$; ****$p < 0.0001$; n.s., not significant), with exact p values reported where shown. Thresholds are expressed in dB SPL. *Abbreviations: SPL, sound pressure level.*

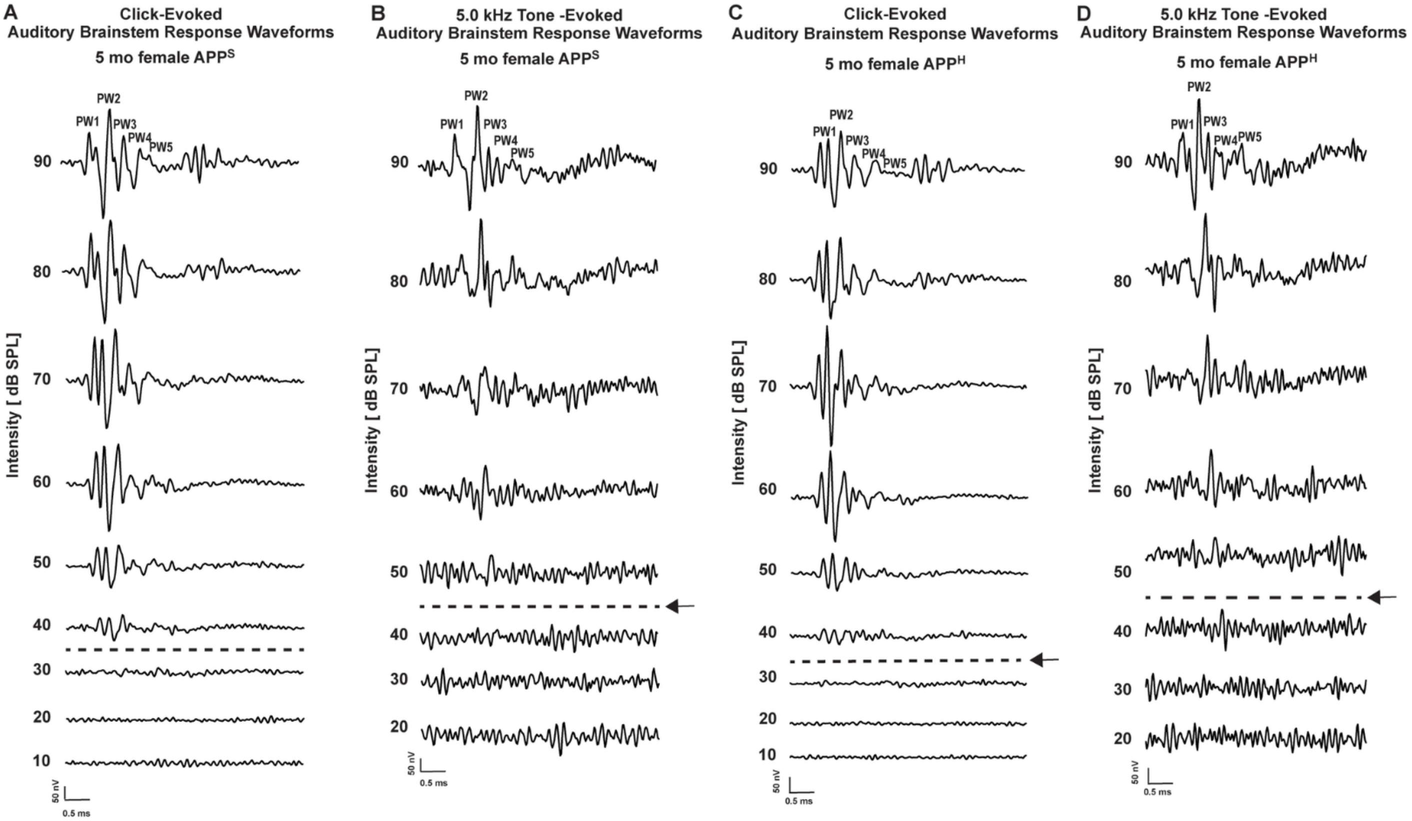


**Supplemental Figure S2:** *Trial-averaged auditory brainstem response (ABR) traces elicited by clicks or tones.* Click-evoked ABRs across sound levels were used to initially determine intact auditory function. Tone-Evoked ABRs across sound levels were used to determine hearing thresholds. **(A-C)**. A representative click-evoked ABR waveforms recorded from a 5-month-old female *App*$^{s}$ and *App*$^{H}$ rats across a range of sound intensities (90–10 dB SPL; in -10 dB steps; traces show 0–10 ms from stimulus onset). The hearing threshold was determined as 40 dB SPL, the lowest sound level that elicits a characteristic ABR waveform (indicated by the dashed line and arrow). **(B-D)** Tone-evoked ABR waveforms recorded from the same animals in response to 5.0 kHz tone bursts presented at intensities ranging from 90 to 20 dB SPL (in -10 dB SPL steps). The tone-evoked threshold was defined as the lowest sound intensity eliciting a detectable response, which here is shown as 50 dB SPL (indicated by the dashed line and arrow). Waveform traces at each sound level represent 512 trial-averaged responses. *Abbreviations: SPL, sound pressure level.*

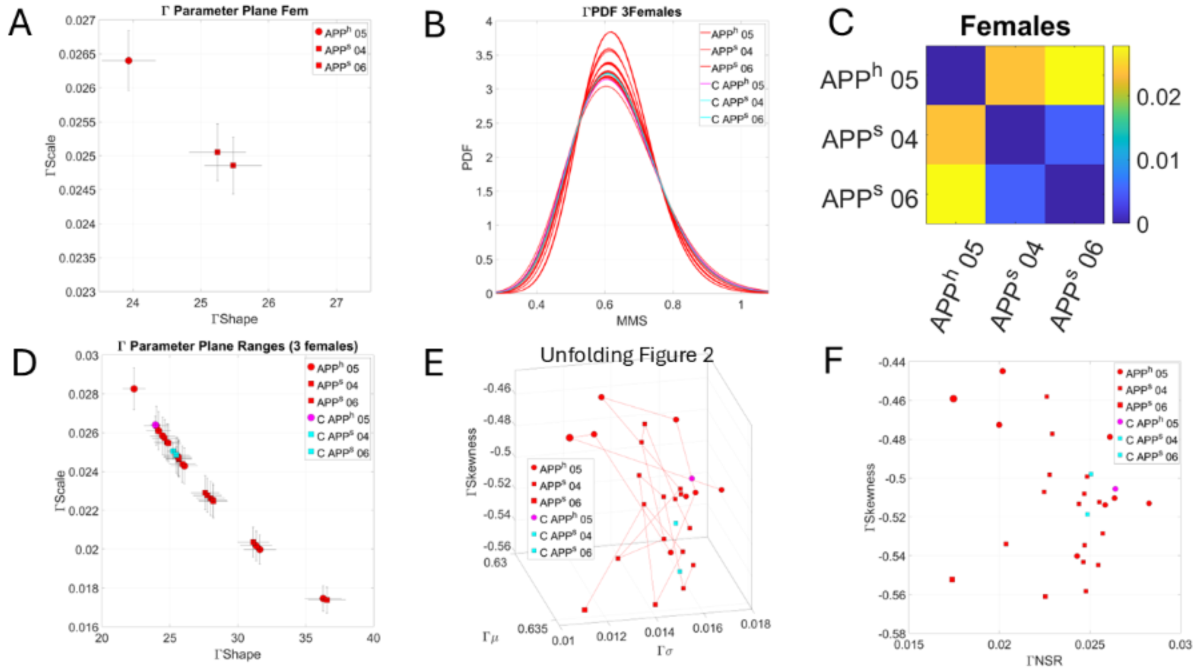


**Supplementary Figure S3**. Unfolding Female data **FIGURE 2**. **(A)** Original **FIGURE 2** plot of the Gamma parameter plane with 3 female rats localized as empirically estimated distribution points with 95% confidence intervals for each of the Gamma shape and Gamma scale parameters. **(B)** Continuous Gamma family of probability distribution functions empirically estimated from the data unfolded second-by-second in 5-second-long blocks for each animal, with 50 % overlapping sliding windows. Red PDFs are the unfolded points while cyan and magenta PDFs represent the centroids of the scatter corresponding to each animal in panel A. **(C)** The Earth Mover's Distance as a proper similarity metric to measure distances between distributions on our Gamma parameter plane. Values of the EMD are represented as a pairwise colormap whereby blue represents smaller distances (more similar distributions) vs. yellow colors representing larger distances (dissimilar distributions). Color bar denotes a range of values. App 04 and 06 mutants are closer in probability space and farther from App 05 humanized **(D)** Representation of the PDFs on the Gamma parameter plane as in (A) for all 24 (8 PDFs x 3 animals) obtained with the sliding window (1 second long) that allowed for tight 95% CI in the Gamma PDF maximum likelihood estimation. **(E)** Stochastic trajectories second-by-second, represented on the Gamma moments parameter space spanned by the empirically estimated Gamma mean (x-axis), Gamma variance (y-axis) and Gamma skewness (z-axis). The fourth moment, the kurtosis value is proportional to the size of the marker, with larger markers representing more kurtotic distributions. As in (D) we plot the centroids in (A) using magenta and cyan colors to represent the mutant and humanized rats. **(F)** Parameter plane spanned by the Gamma NSR on the x-axis (summarizing the Gamma var/Gamma mean) and the Gamma skewness on the y-axis. This projection helps us visualize the relationship of the centroid PDFs in A and the scatter describing the family of PDFs of the female cohort.

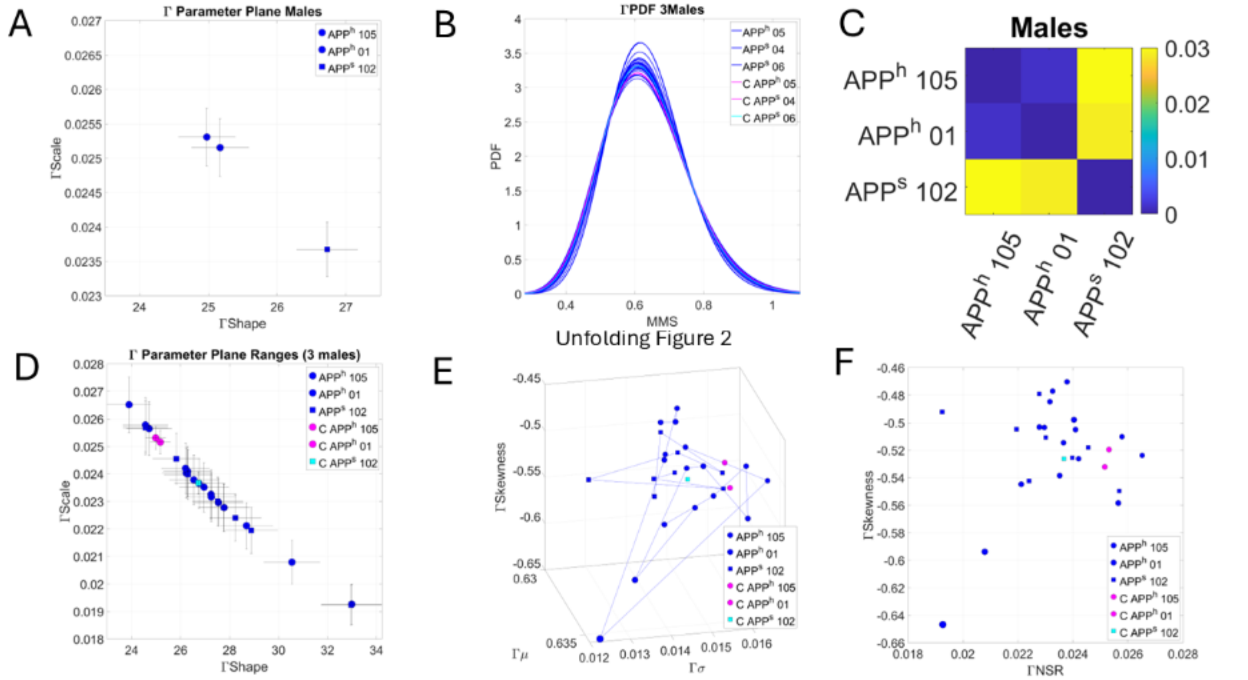


**Supplementary Figure S4.** Similarly to the above figure, the male rats in FIGURE 2 consistently show differences between the empirically estimated Gamma families **(A-B)** obtained from the responses of mutants and humanized rats. These differences are objectively quantified through the EMD **(C)** and unfolded on the Gamma moments space **(D-E)** and the projection space **(F)**.

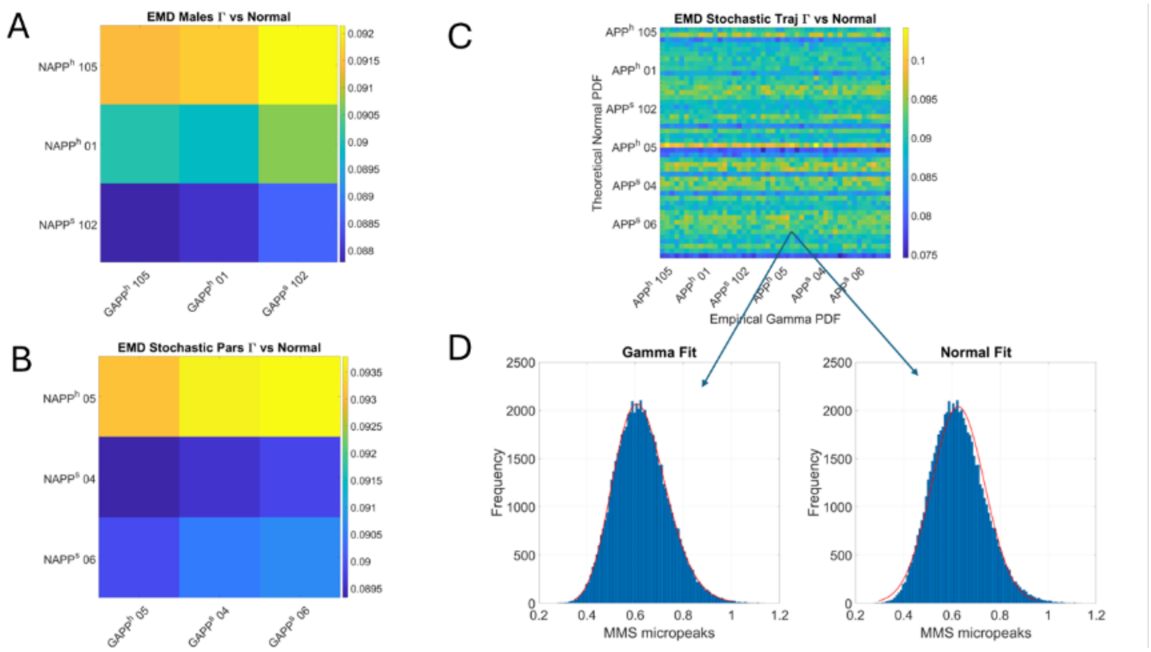


**Supplementary Figure S5.** Quantification of differences in families of probability distribution functions empirically estimated from the data rather than theoretically imposed. **(A-B)** Males and females empirically estimated Gamma PDFs of the centroids in **FIGURE 2** of the main text are compared against the assumed theoretical normal distribution. Differences are represented through the color map matrix of comparisons across each rat. Notice the range of differences in the color bar indicating non-zero values denoting a departure from (often assumed) theoretical normality when obtaining grand averages. **(C)** Unfolding the stochastic trajectories gives continuous Gamma families of 24 points for each of the males and females cohorts. The matrix with entries representing values of the EMD comparing the theoretical Normal vs. the empirically obtained Gamma families, further shows the differences for the second-by-second sampling with 50% sliding window overlap. **(D)** Representative fitting of the histograms of micro peaks (micro movement spikes peaks) shows the lack of fitting of the imposed Normal distribution.

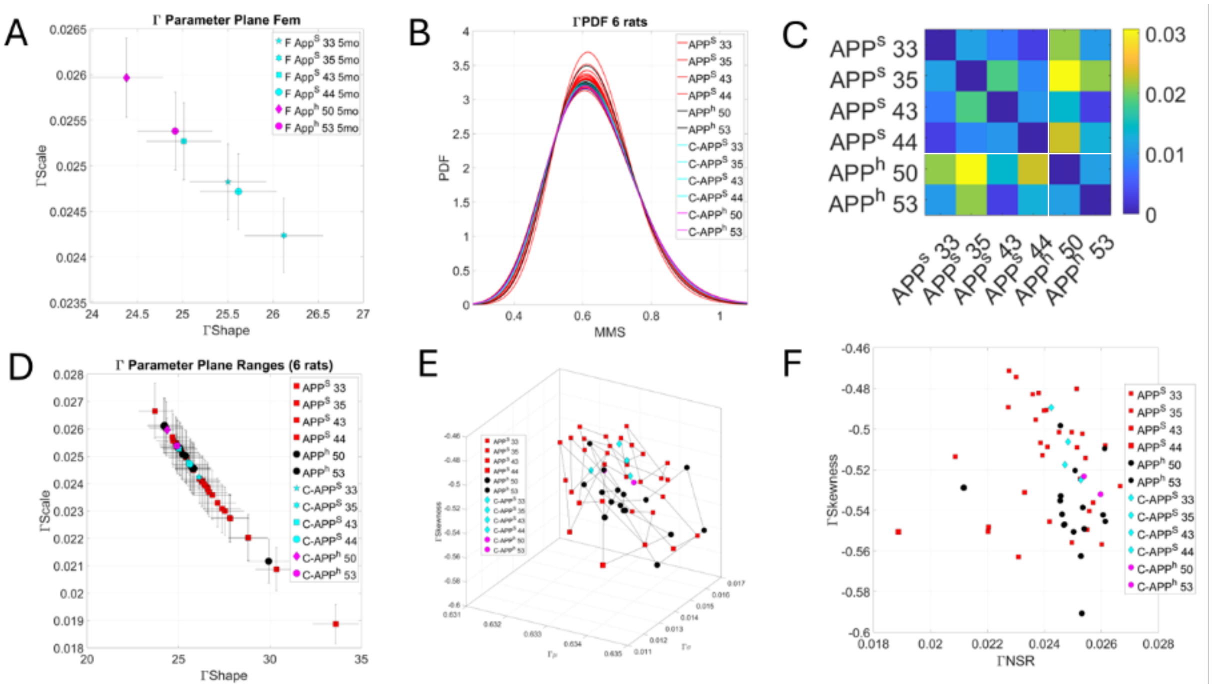


**Supplementary Figure S6.** Unfolding stochastic trajectories on the Gamma parameter plane derived from responses of younger female animals from **FIGURE 3** in the main text. **(A)** Empirically estimated continuous Gamma family of probability distribution functions (PDFs) localizing each animal as a pair of (shape, scale) values with 95% confidence intervals. These are the estimated PDFs of the full response. **(B)** The Gamma family across this cohort obtained from one-second-long sliding window with 50% overlap. Points are plotted as Gamma PDFs along with the values corresponding to panel A. **(C)** Proper similarity metric EMD to quantify, pairwise, the distances in probability space between each of the Gamma PDFs in (A). Distances are in correspondence with localized parameters on the Gamma plane. (D) Unfolded stochastic trajectories on the Gamma plane. **(E)** Corresponding unfolded trajectories on the Gamma moments space. **(F)** Projection of parameter on the Gamma NSR vs. the Gamma Skewness dimensions across all trajectories and including the centroid points as well in (A).

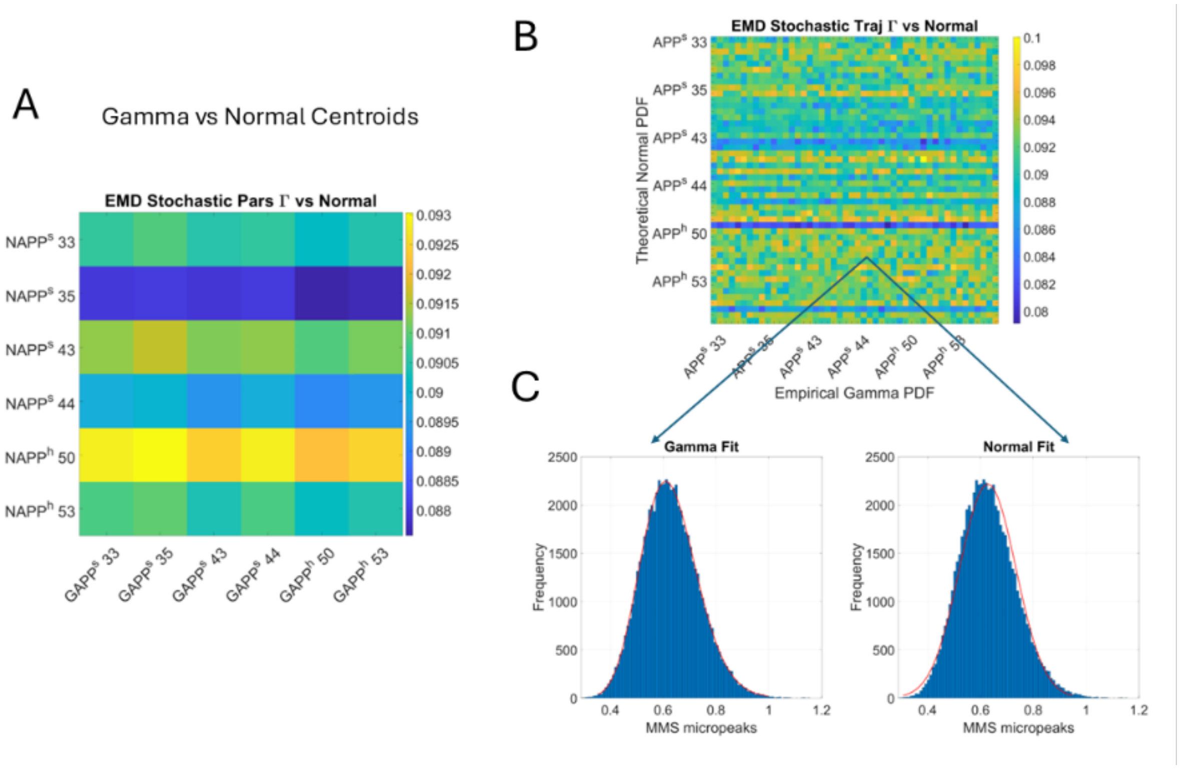


**Supplementary Figure S7.** Quantification of statistical differences according to appropriate similarity metric (the EMD) comparing the theoretical normal to the empirically estimated Gamma family. **(A)** Pairwise comparisons for each animal in FIGURE 3 of the main text are displayed as a color-coded matrix. Entries are the EMD value (range on the color bar), all different from zero, denoting the departure from normality for each point in (A). **(B)** Unfolding trajectories second-by-second for each animal and comparing the assumed Theoretical Normal vs the empirically estimated Gamma family. **(C)** Representative fitted distributions to the MMS micro peaks of one data point in B, visualizing the misfit of the theoretical normal.

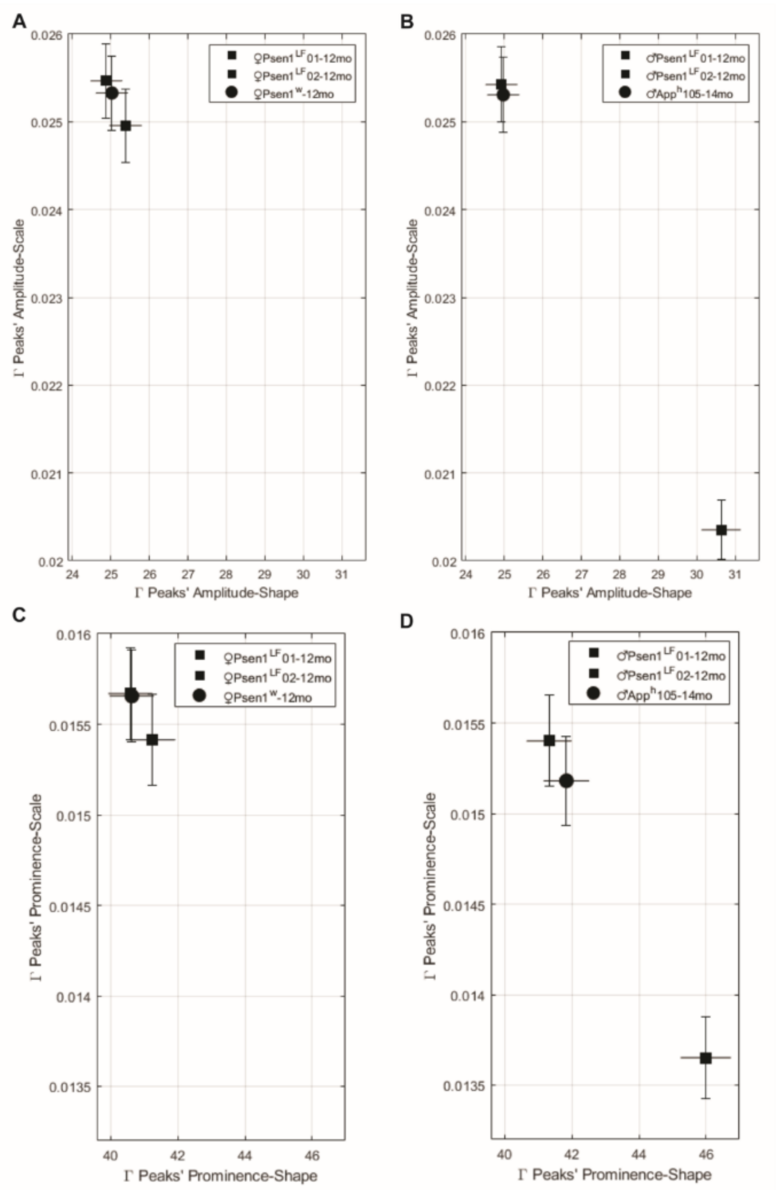


**Supplemental Figure S8:** *Multiparametric auditory brainstem response (ABR) micro-peak feature extraction on distinct AD risk genotypes may reveal mutation-specific signatures, while preserving within-genotype consistency.* (A- B) In contrast to the *App*$^{S}$ risk mutation (see **FIGURE 2**), fluctuations in the amplitudes of ABR "micro-peaks" fail to separate both female (left) and male (right) *Psen1*$^{LF}$ rats (square icons) from humanized control rats (*Psen1*$^{w}$, circle icons, right; *Psen1*$^{w}$ */App*$^{H}$, circle icons, left) in the same 2-dimensional state space (i.e., gamma shape and scale). (C-D) Alternate variability measures of micro-peak prominence also show overlap of *Psen1*$^{LF}$ rats (square icons) with *Psen1*$^{w}$ (circle icons) in an alternate state space in both sexes. That the PSEN1 mutation does not appear to separate in state space from healthy controls (e.g., *Psen1*$^{w}$ */App*$^{H}$ rats) suggests a different neurophysiological phenotype arising from APP vs. PSEN1 mutations. *Note the labelled axes in panels A-B vs. C-D as they represent different features and thus state space.*

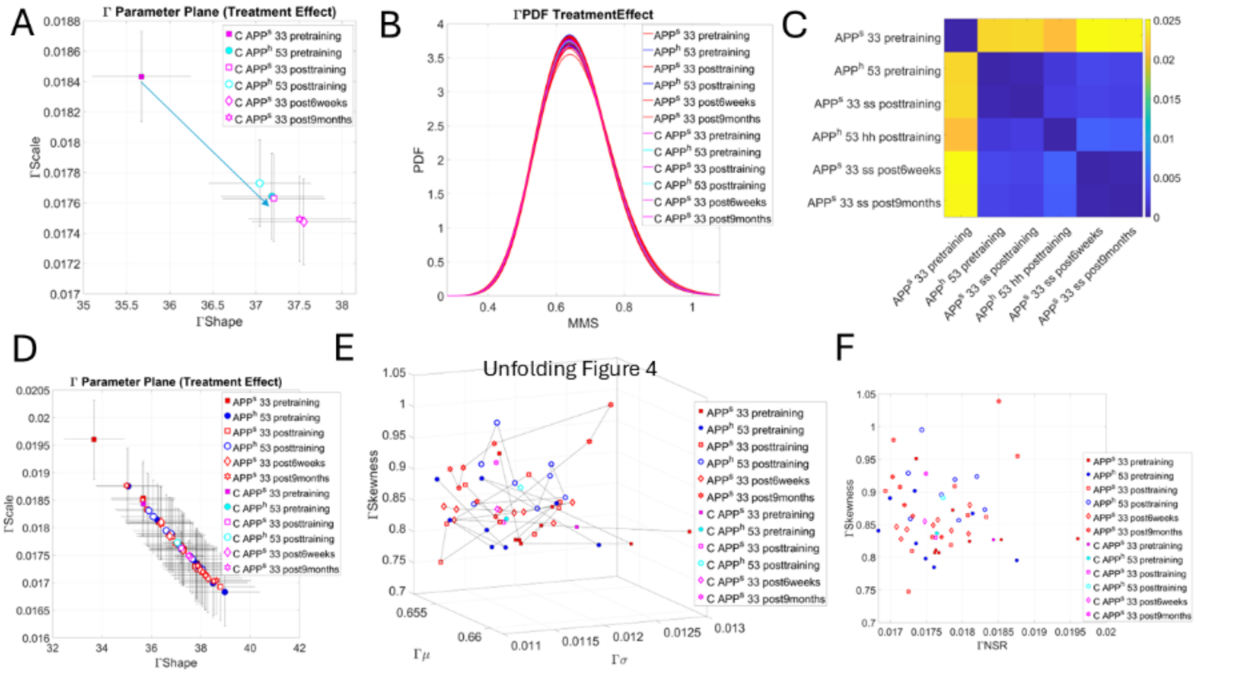


**Supplemental Figure S9.** Effects of training on responses time. **(A)** Empirically estimated distributions of the inter-peaks interval times from **FIGURE 4**, plotted as points on the Gamma parameter plane spanned by the shape and scale. Arrow indicates largest stochastic shift between rat *App*$^s$ 33 pre-training and post training responses. A decrease in the Gamma NSR (scale) given by the Gamma variance/Gamma mean ratio. **(B)** Family of continuous Gamma distributions from unfolding each block into corresponding signatures over one-second-long sliding windows with 50% overlap. Gamma PDFs from the centroids from panel (A) are also included. **(C)** EMD quantifying the PDF distances represented as a colormap obtained pairwise between the PDFs in (A). **(D)** Unfolding the points in (A) provides the span of the ranges for each animal. **(E)** Localizing the stochastic trajectories on the Gamma moments space along with the points in (A) denoted C (centroids). **(F)** Projection of the PDF points on the parameter plane spanned by the Gamma NSR and the Gamma skewness.

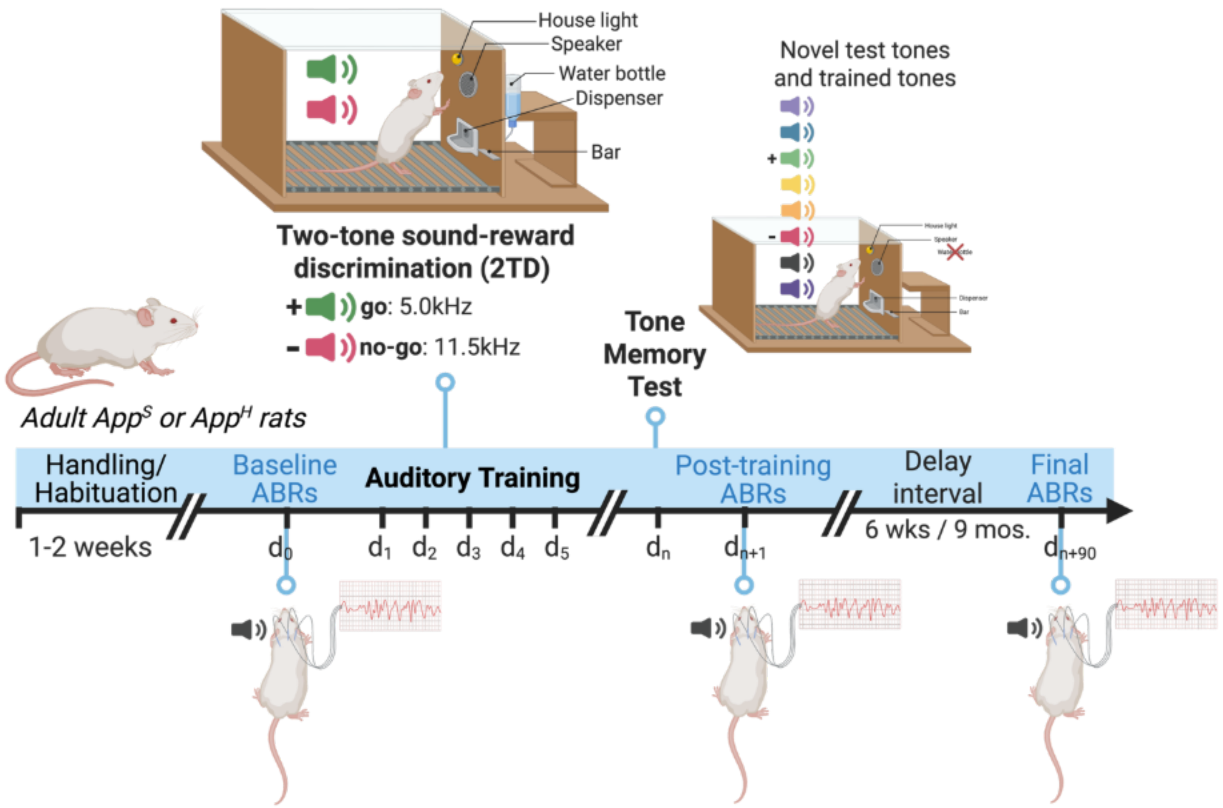


**Supplemental Figure S10:** *Experimental timeline*. Behavioral training was in an auditory cognitive task for sound-reward learning in parallel with electrophysiology in *App^S* and *App^H* rats. All rats were handled to habituate animals to experimenters and the lab space prior to beginning the experimental testing and training procedures. A subset of animals with ABR recordings underwent sound-guided cognitive training in a two-tone discrimination task (2TD), followed by a memory test. The 2TD task requires rats to bar press to a rewarded "go" tone (5.0 kHz; 75 dB SPL) for a water reward, and to omit responding to an unrewarded "no-go" tone (11.5 kHz; 75 dB SPL) to avoid time-outs that extend the time to the next trial. A memory test occurred 24 hrs after the final training session and included eight pure tones of different frequencies (including the "go" and "no-go" frequencies). Bar press numbers were recorded and analyzed for each test tone separately to determine the success of auditory memory formation for the rewarded "go" tone after training. Tone-evoked ABRs were recorded in trained animals at three timepoints: prior to the first 2TD training session ("baseline" pre-training timepoint), 24 hrs after the memory test (post-training timepoint), and after a delay (6 weeks later and 9 months later) to track the persistence of training intervention-induced changes in auditory neurophysiological signals longitudinally.

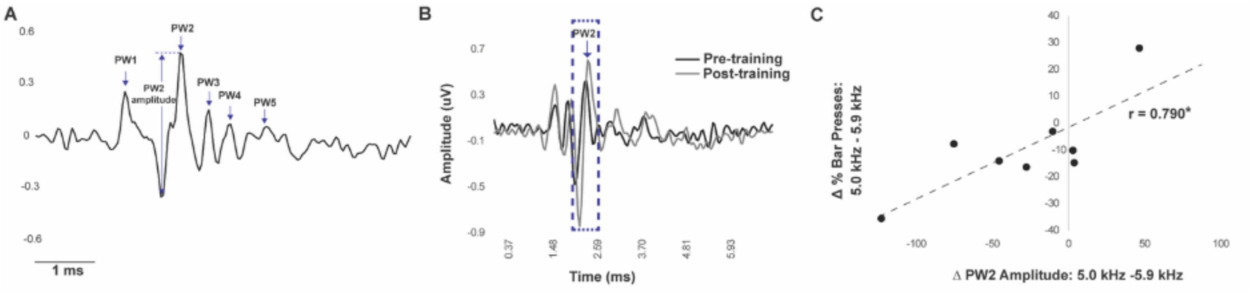


**Supplemental Figure S11:** *Auditory brainstem response (ABR) waves relate to successful memory formation.* **(A)** Representative sound-evoked ABR waveform that can be recorded from a rat, similarly to humans, that shows identified 5 positive waves, PW1–PW5. The amplitude of a wave is measured from the maximum peak of the wave to the minimum of the preceding trough in time (a voltage). For example, PW2 amplitude is defined as the voltage difference between the preceding trough minimum and the PW2 peak maximum. **(B)** Representative ABR traces evoked by the rewarded tone (5.0 kHz) at before ("pre-training," black) and after ("post-training," gray) two tone reward training time points. There was an increase in the PW2 amplitude in ABRs evoked by a rewarded signal tone from pre- to post- two-tone reward training. Dashed boxes indicate PW2. (middle). **(C)** There is a significant positive correlation between differences in PW2 amplitude ABRs evoked by the 5.0 kHz signal tone vs. a nearby neighbor tone (5.9 kHz) and the difference in percentage of responses made to the 5.0 kHz tone vs 5.9 kHz during the memory test. Dots indicate individual data points. Greater amplitude differences predict a greater difference in the percentage of responses allocated to the signal tone frequency (*Two-tailed, Pearson's R = 0.790; p=0.0197).

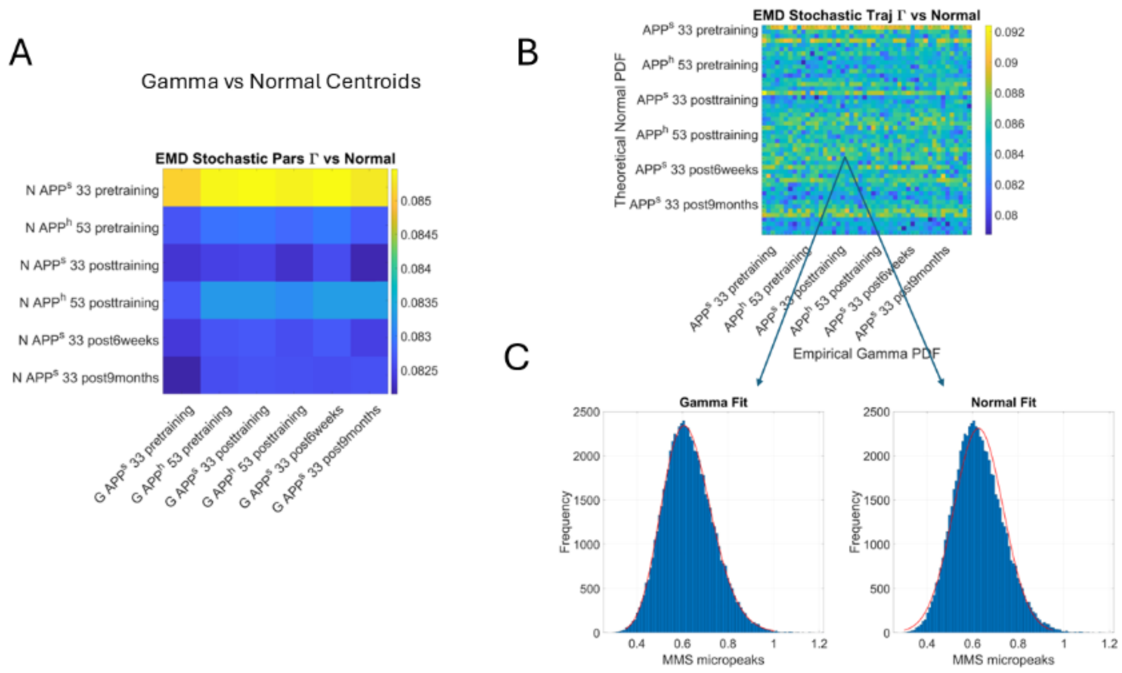


**Supplementary Figure S12.** Quantification of disparity between the theoretically assumed Normal distribution and the empirically estimated Gamma distribution across the sample in **FIGURE 4.** **(A)** Pairwise EMD between normal and gamma fits for the centroids. **(B)** Pairwise EMD between normal and gamma fits for each second-long window for each animal. Notice that in both the centroids and unfolded data the minimum value of the EMD is not zero. **(C)** Representative distributions with the fit (red) PDF showing the inadequacy of the Normal distribution fit on the histograms of the MMS micro peaks.

### SUPPLEMENTARY TABLES

**Supplementary Table S1.** This table presents individual click- and tone- evoked ABR hearing thresholds.

| | | | Individual ABR Thresholds (dB SPL) by Frequency | | | | |
|---|---|---|---|---|---|---|---|
| Subject | Sex | Age (Months) | Click | *pre*-training 5 kHz | *pre*-training 11.5 kHz | *post*-training 5kHz | *post*-training 11.5 kHz |
| App$^{s}$33 | F | 5 | 40 | 60 | 70 | 60 | 50 |
| App$^{s}$35 | F | 5 | 45 | 70 | 60 | | |
| App$^{s}$43 | F | 5 | 35 | 60 | 50 | | |
| App$^{s}$44 | F | 5 | 40 | 50 | 50 | | |
| App$^{h}$50 | F | 5 | 40 | 50 | 50 | | |
| App$^{h}$53 | F | 5 | 40 | 50 | 50 | 50 | 50 |
| App$^{s}$04 | F | 18 | 45 | 60 | | | |
| App$^{h}$05 | F | 18 | 45 | 70 | | | |
| App$^{s}$06 | F | 17 | 40 | 70 | | | |
| App$^{h}$01 | M | 26 | 50 | 80 | | | |
| App$^{s}$102 | M | 13 | 50 | 60 | | | |
| App$^{h}$105 | M | 14 | 50 | 80 | | | |
| PSEN1$^{w}$ | F | 12 | 50 | 70 | | | |
| PSEN$^{LF}$-01 | F | 12 | 40 | 70 | | | |
| PSEN$^{LF}$-02 | F | 12 | 50 | 50 | | | |
| PSEN$^{LF}$-01 | M | 12 | 50 | 70 | | | |
| PSEN$^{LF}$-02 | M | 12 | 50 | 80 | | | |

**Supplementary Table S2.** This table presents individual pre- and post-training 5.0 kHz tone evoked ABR PW2 amplitude and latency values.

| | | | **Individual Sound Evoked ABR PW2 Analysis** | | | |
|---|---|---|---|---|---|---|
| Subject | Sex | Age (Months) | *pre*-training 5.0 kHz PW2 Amplitude (nV) | *pre*-training 5.0 kHz PW2 Latency (ms) | *post*-training 5kHz PW2 Amplitude (nV) | *post*-training 5.0 kHz PW2 Latency (ms) |
| $App^s$33 | F | 5 | 209.72 | 2.42 | 184.19 | 2.46 |
| $App^s$35 | F | 5 | 154.13 | 2.46 | - | - |
| $App^s$43 | F | 5 | 313.16 | 2.62 | - | - |
| $App^s$44 | F | 5 | 291.21 | 2.62 | - | - |
| $App^h$50 | F | 5 | 300.54 | 2.58 | - | - |
| $App^h$53 | F | 5 | 281.29 | 2.75 | 327.41 | 2.66 |
| $App^s$04 | F | 18 | 162.97 | 2.70 | - | - |
| $App^h$05 | F | 18 | 70.21 | 2.79 | - | - |
| $App^s$06 | F | 17 | 319.95 | 2.87 | - | - |
| $App^h$01 | M | 26 | 109.49 | 2.42 | - | - |
| $App^s$102 | M | 13 | 186.09 | 2.58 | - | - |
| $App^h$105 | M | 14 | 117.39 | 2.79 | - | - |
| $PSEN1^w$ | F | 12 | 277.53 | 2.75 | - | - |
| $PSEN^{LF}$-01 | F | 12 | 310.69 | 2.66 | - | - |
| $PSEN^{LF}$-02 | F | 12 | 217.83 | 2.62 | - | - |
| $PSEN^{LF}$-01 | M | 12 | 192.48 | 2.50 | - | - |
| $PSEN^{LF}$-02 | M | 12 | 126.26 | 2.50 | - | - |

**Supplemental Table S3.** The table below presents the PW2 amplitude (M ± SE) from the comparisons used in each figure. Sample sizes are in parentheses.

| | | | ***App<sup>S</sup>*** | | ***App<sup>H</sup>*** | |
|---|---|---|---|---|---|---|
| | | | ***Amplitude (nV)*** | ***Latency (ms)*** | ***Amplitude (nV)*** | ***Latency (ms)*** |
| **Figure 2A** | PW2 | 5.0 kHz | 241.46 ± 78.49 (n=2) | 2.78 ± 0.08 | 70.21 ± 0 (n=1) | 2.78 ± 0 |
| **Figure 2B** | PW2 | 5.0 kHz | 186.09 ± 0 (n=1) | 2.58 ± 0 | 113.43 ± 3.94 (n=2) | 2.60 ± 0.18 |
| **Figure 3** | PW2 | 5.0 kHz | 242.05 ± 36.79 (n=4) | 2.53 ± 0.05 | 290.91 ± 9.62 (n=2) | 2.66 ± 0.08 |
| **Figure 4** | PW2 | pre- vs. post | 245.50 ± 35.78 (n=2) | 2.58 ± 0.16 | 255.79 ± 71.61 (n=2) | 2.56 ± 0.10 |

| | | | **Psen1<sup>w</sup>** | | ***Psen1<sup>LF</sup>*** | | ***App<sup>H</sup>*** | |
|---|---|---|---|---|---|---|---|---|
| | | | ***Amplitude (nV)*** | ***Latency (ms)*** | ***Amplitude (nV)*** | ***Latency (ms)*** | ***Amplitude (nV)*** | ***Latency (ms)*** |
| **Supplemental Figure 1A** | PW2 | 5.0 kHz | 277.52 ± 0 (n=1) | 2.74 ± 0 | 264.26 ± 46.42 (n=2) | 2.78 ± 0 | - | - |
| **Supplemental Figure 1B** | PW2 | 5.0 kHz | - | - | 159.36 ± 33.10 (n=2) | 2.50 ± 0 | 117.38 ± 0 (n=1) | 2.78 ± 0 |

**Supplemental Table S4.** The table below presents the number of animals included in each comparison.

| Group | Female | | Male | |
|---|---|---|---|---|
| | Young rats (5 months old) | Aged (>12 months old) | Young rats (5 months old) | Aged (>12 months old) |
| *App*$^{S}$ | n = 4$^{\#}$ | n = 2$^{\$}$ | n/a | n = 1* |
| *App*$^{H}$ | n = 2$^{\#}$ | n = 1$^{\$}$ | n/a | n = 2* |
| *Psen1*$^{LF}$ | n/a | n = 2$^{@}$ | n/a | n = 2$^{@}$ |
| *Psen1*$^{W}$ | n/a | n = 1$^{@}$ | n/a | n/a |

*denotes the groups and numbers of animals included in the analysis shown in Figure 2B.

$denotes the number of animals included in the analysis shown in Figure 2A.

#denotes the groups and numbers of animals included in the comparisons shown in Figure 3.

@denotes the groups and numbers of animals included in the comparisons shown in Supplemental Figure 1.

*Notes:* One aged *App*$^{H}$ rat (*) was also included in the comparisons shown in Supplemental Figure 1. Figure 4 includes n = 2 rats (#) (1 *App*$^{S}$ and 1 *App*$^{H}$) used for the pairwise comparisons (*pre- vs post- training*)
